# Marine cold seep ANME-2/SRB consortia produce their lipid biomass from inorganic carbon

**DOI:** 10.1101/2025.08.02.667857

**Authors:** Lennart Stock, Gunter Wegener, Yueqing Wang, Yannick Zander, Marcus Elvert

## Abstract

In cold seeps, anaerobic methanotrophic archaea (ANME) and sulfate-reducing bacteria (SRB) oxidize methane to inorganic carbon (IC) coupled to sulfate reduction. While catabolic pathways are well resolved, carbon flow into biomass remains unclear. We conducted lipid stable isotope probing (lipid-SIP) experiments with Astoria Canyon sediments dominated by ANME-2/SRB consortia and incubated samples with either ^13^C-labeled methane (^13^CH_4_) or dissolved IC (DI^13^C). Lipid-specific δ^13^C analysis showed higher ^13^C incorporation from DI^13^C than from ^13^CH_4_. After 30 days, δ^13^C values were up to +417‰ in SRB-specific fatty acids (e.g., C_16:1ω5c_, cyC_17:0ω5,6_) and +126‰ in ANME-2-specific isoprenoid lipids (e.g., archaeol, crocetane). Based on these values, we calculated carbon assimilation rates and found that both partners primarily assimilate IC. Remarkedly, IC assimilation in SRB lipids was eight times higher than in ANME lipids, suggesting that ANME either rely on fixation of internally generated CO_2_ from ^13^C-label-free methane before it can be exchanged with the environment or they utilize an additional, yet unknown carbon source for lipid biosynthesis. By examining the step-wise ^13^C-enrichment of ANME- and SRB-derived lipids, we further delineate biosynthetic pathways for archaeal and bacterial diether lipid formation and highlight crocetane as a bilayer-modulating isoprenoid hydrocarbon potentially affecting membrane fluidity and proton permeability.

**Graphical abstract:** 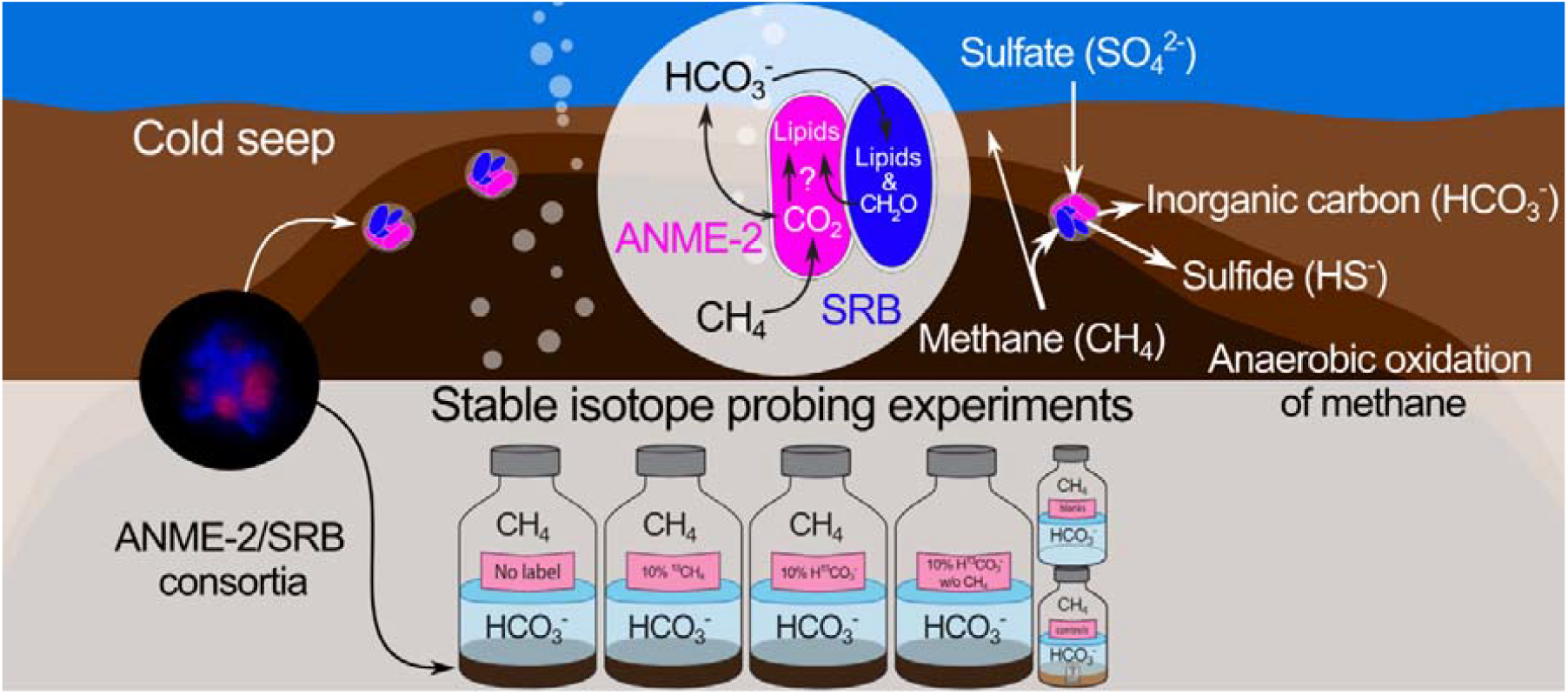

Lipid-SIP experiments in Astoria Canyon sediments revealed that both ANME-2 and SRB primarily assimilate inorganic carbon (IC), not methane, into biomass. SRB-specific lipids showed eightfold higher I^13^C-assimilation than ANME lipids, suggesting SRB directly assimilate environmental IC, while ANME rely on internally produced CO_2_ or alternative carbon sources, potential provided by SRBs.

## Introduction

The anaerobic oxidation of methane (AOM) with sulfate as electron acceptor is a major methane sink, consuming approximately 90 % of the methane produced in marine sediments (Hinrichs and Boetius, 2002; Knittel and Boetius, 2009; Reeburgh, 2007). AOM is mediated by syntrophic consortia composed of anaerobic methanotrophic archaea (ANME), which oxidize methane to inorganic carbon (IC), and sulfate-reducing bacteria (SRB) that use reduction equivalents of AOM to reduce sulfate to sulfide (Orphan et al., 2001). The overall redox reaction for AOM coupled to sulfate reduction (SR) can be summarized as CH_4_ + SO_4_^2-^ → HCO_3_^−^ + HS^−^ + H_2_O, with a δG at under in situ conditions of −20 to −40 kJ mol^−1^ (Knittel and Boetius, 2009; Larowe et al., 2008; Thauer, 2011).

ANME archaea comprise three polyphyletic linages within the phylum Halobacteriota. ANME-1 (*Ca*. Methanophagales) forms a distinct order dominating deep sulfate methane transition zones (SMTZ) and hydrothermal and hypersaline environments (Knittel et al., 2019; Ruff et al., 2015). ANME-3 (*Ca*. Methanovorans) is a genus within the family Methanosarcinacea, and is closely related to Methanococcoides. This uncultured ANME group is often found at mud volcanoes (Lazar et al., 2011; Niemann et al., 2006). ANME-2 dominates most cold seep environments, especially those with high AOM activity (e.g., Niemann et al., 2006; Omoregie et al., 2008; Treude et al., 2003). This linage includes several subgroups, including ANME-2a, −2b (*Ca*. Methanocomedens, *Ca*. Methanomarinus), −2c (*Ca*. Methanogaster) (Chadwick et al., 2021; Knittel and Boetius, 2009; Wegener et al., 2022) and the terrestrial ANME-2d clade (*Ca*. Methanoperedens, Haroon et al., 2013).

Metagenomic and metatranscriptomic studies revealed that all ANME linages oxidize methane via a C_1_-pathway resembling reverse methanogenesis (Chadwick et al., 2022; Hallam et al., 2004; Meyerdierks et al., 2010; Thauer, 2011; Wang et al., 2014). Most marine ANME grow in syntrophy with SRB of the Desulfobacterota phylum, as demonstrated by fluorescence in situ hybridization (FISH, e.g., Boetius et al., 2000; Metcalfe et al., 2021; Orphan et al., 2001; Osorio-Rodriguez et al., 2023). In cold seep environments, SRB typically belong to the Desulfococcus/Desulfosarcina (DSS) clade (e.g., Boetius et al., 2000; Orphan et al., 2001) and derive the reducing equivalents from ANME, utilizing the reducing power for sulfate reduction (McGlynn et al., 2015; Wegener et al., 2015). Recent studies suggest that the consortia members perform direct interspecies electron transfer mediated by cytochrome-based nanowires (Chadwick et al., 2022; McGlynn et al., 2015; Wegener et al., 2015). Metagenomic analysis indicate that most SRB encode the Wood–Ljungdahl pathway, suggesting that the partner bacteria are autotrophs (Hügler and Sievert, 2011; Skennerton et al., 2017). Similarly, ANME encode an often slightly modified archaeal version of this pathway (Chadwick et al., 2022). However, in ANME the C1-branch of its Wood-Ljungdahl pathway is central to reverse methanogenesis pathways, which enable the oxidation of methane to CO_2_ (Thauer, 2011).

Both, ANME and the SRB partner produce characteristic ^13^C-depleted membrane lipids, indicating the incorporation of methane-derived carbon. ANME-2 archaea typically produce the glycerol dialkyl diethers sn-2 hydroxyarchaeaol (sn2-OH-Ar) and archaeol, while ANME-1 archaea produce mainly glycerol dialkyl glycerol tetraethers (GDGTs), all with δ^13^C values ranging from of −70‰ to −130‰ relative to Vienna Pee Dee Belemnite (VPDB) (e.g., Elvert et al., 2005, 1999; Hinrichs et al., 2000; Niemann and Elvert, 2008; Orphan et al., 2002). These are among the most ^13^C-depleted biogenic compounds known and are substantially lower than those of other autotrophic microorganisms, such as phototrophic algae which produce lipids and biomass with δ^13^C values between −18‰ to −36‰ (Goericke and Fry, 1994; Henderson et al., 2024). The dominant SRB lipids in AOM consortia include fatty acids such as C_16:1ω5c_ and cyC_17:0ω5,6_, with δ^13^C values between –60‰ and –100‰ (e.g., Blumenberg et al., 2004; Elvert et al., 2003). The variability in lipid δ^13^C values of ANME and SRB reflects the wide carbon isotope range of environmental methane (–10‰ to –100‰), depending on whether it is of thermogenic or biogenic origin (Claypool and Kaplan, 1974; Whiticar, 1999). Because AOM generates ^13^C-depleted dissolved inorganic carbon (DIC), the question remains whether consortia members assimilate methane, DIC, both, or additional carbon sources.

To resolve this question, multiple studies have repeatedly applied lipid stable isotope probing (lipid-SIP; Boschker and Middelburg, 2002; Wegener et al., 2016a) to track the assimilation of ^13^C-labeled carbon substrates into microbial lipids of AOM active cold seep sediments, microbial mats, and enrichment cultures (Bertram et al., 2013; Blumenberg et al., 2005; Jagersma et al., 2009; Kellermann et al., 2012; Wegener et al., 2016b, 2012, 2008). However, this often led to different conclusions. Early studies used only ^13^CH_4_ methane (Blumenberg et al., 2005; Jagersma et al., 2009), whereas subsequent work tested also DI^13^C as potential carbon substrate (Kellermann et al., 2012; Wegener et al., 2016b, 2012, 2008). Some studies, particularly on ANME-2d archaea, which perform nitrate-dependent AOM, utilize both methane and IC, hence suggesting a mixotrophic growth (Kurth et al., 2019). In contrast, ANME-1 archaea appear to preferentially assimilate IC, indicating chemoautotrophy as the dominant mode (Kellermann et al., 2012; Treude et al., 2007; Wegener et al., 2016b). In case of ANME-2, however, the carbon assimilation is less clear, as a conclusive distinction between methane and inorganic carbon sources is not established yet (Wegener et al., 2008).

In this study, we conducted lipid-SIP experiments with cold seep sediments from the Astoria Canyon off the Oregon coast, which are dominated by marine ANME-2/SRB consortia. Under AOM favorable conditions, we incubated the samples with DI^13^C, with and without CH_4_, as well as with ^13^CH_4_ and DIC. This design allowed us to quantify AOM-dependent carbon assimilation and lipid turnover from either IC or methane. Our results provide new insights into the metabolic strategies of ANME-2, in particular its ability to produce its biomass from IC. In addition, we tracked the time course of ^13^C-assimilation into archaeal and bacterial lipids, which sheds light on lipid biosynthetic pathways and supports the hypothesis that archaea synthesize isoprenoid hydrocarbons as membrane lipid modulators.

## Material and Methods

### Site description and sample collection

The Astoria Canyon, located on the northern Cascadia margin, hosts numerous cold seep sites, reflecting the widespread occurrence of such features across the entire Cascadian margin (Baumberger et al., 2018; Merle et al., 2021). During the R/V Atlantis cruise AT50-14 in August 2023, we collected sediment cores from the cold seep site “H1867” in Astoria Canyon, Oregon, USA (lat.: 46.240784°N, long.: −124.602954°W, water depth: 760 m). Geochemical characterization of the same cold seep system can be freely accessed through the Biological and Chemical Oceanography Data Management Office (BCO-DMO) repository (www.bco-dmo.org/dataset/959765) (Lalk et al., 2025).The location was previously found during E/V Nautilus cruise NA128 with the ROV Hercules (Dive H1867; Wishnak, 2022). The first 20 cm of the push core were pooled in glass bottles and were kept under anoxic conditions until further preparation of the experiment in the laboratory.

### Incubation of sediment slurries

A sediment slurry was prepared with 200 g of cold seep sediment and 500 mL artificial anoxic seawater medium (Laso-Pérez et al., 2018) with 25 mM magnesium sulfate and 30 mM sodium bicarbonate (i.e., DIC; Merck). These slurries were then evenly distributed into six 250-mL serum vials and filled up to 100 mL with the anoxic medium. The vials were incubated with CH_4_ as energy source. Rates of methane-dependent sulfide production were monitored using an established colorimetric assay (Cord-Ruwisch, 1985; Laso-Pérez et al., 2018). All samples showed instant sulfide production (>5 mM in 30 days). For the SIP experiment, all slurries were pooled and supplied with new medium with 27 mM DIC. This slurry was distributed in 13 250-mL serum vials. Each bottle received 50 mL sediment slurry and 100 mL anaerobic seawater medium, leaving 100 mL gas headspace. Each vial was spiked with either ^13^C-labeled or non-labeled methane or DIC to achieve the following experimental conditions (see Table 1 for details): (1) 30 mM DIC and ∼30 mM CH_4_, (2) 10% labeled 30 mM DIC and ∼30 mM CH_4_, (3) 30 mM DIC and 10% labeled ∼30 mM CH_4_ and (4) 30 mM DIC and no CH_4_. We aimed for 30 mM concentrations and a labeling strength of 10% ^13^C for both DIC and CH_4_ and received ∼5000‰ DIC and ∼10,000‰ CH_4_ isotope background values (Table 1). In addition, blank incubations with about 5 g combusted sea sand were prepared for the treatments (1), (2) and (3), while killed controls, using the original sediments, were set up for the treatments (2) and (3). For the latter two, additions of 10 mL 20% Zink chloride (Merck) solution were used. Treatments containing labeled DIC received 3 mM ≥ 99% of DI^13^C (Sigma-Aldrich), while non-labeled treatments received 3 mM non-labeled DIC to compensate for the missing 3 mM. All treatments containing CH_4_ received 1 bar of CH_4_/CO_2_ (90/10 %v/v) (Air Liquide). Treatments containing labeled CH_4_ received 10 mL of ≥ 99% ^13^CH_4_ (Sigma-Aldrich). For treatments without CH_4_, a gas mixture of N_2_/CO_2_ (90/10 %v/v; Air Liquide) of 1 bar was used.

**Table 1:**
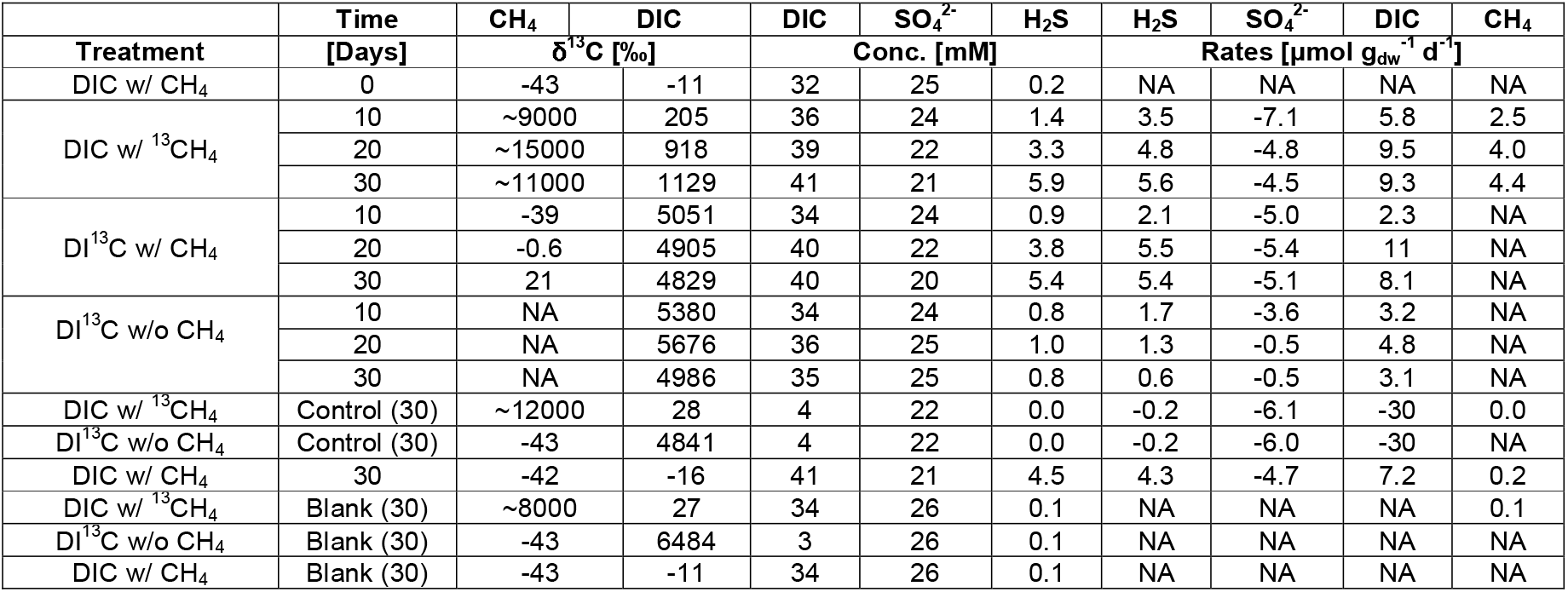
Overview of isotopic compositions and geochemical data of all individual SIP incubation experiments. Listed are δ_13_C values of methane (CH_4_) and dissolved inorganic carbon (DIC), as well as the concentrations of DIC, sulfate (SO_4_^2-^) and hydrogen sulfide (H_2_S). Treatments with “^13^C” stand for a ^13^C labeling of ∼10%. Production and consumption rates of SO_4_ ^2-^, H_2_S, DIC, and CH_4_ are given in µmol g_dw_ ^−1^ d^−1^, NA: not analyzed.

### Methane stable carbon isotope analysis

After completion of the experiment, the headspace methane of all samples was analyzed for stable carbon isotope compositions. In brief, 1 mL headspace gas was transferred into a N_2_-purged 12 mL gas-tight exetainer vial filled with 1 mL concentrated NaOH (18 M, Merck). Vials were stored at room temperature up-side down, to minimize gas escape until measurements. The δ^13^C values of methane were determined using gas chromatography (Trace GC ultra) coupled via a GC combustion III interface to a Delta Plus isotope ratio mass spectrometer (IRMS; all ThermoFinnigan). Gas chromatography involved a Carboxen® 1006 PLOT fused Silica capillary column (30 m x 0.23 mm, Supelco) and helium as carrier gas at a constant flow rate of 3.0 mL min^−1^. 100 µL sample volume was injected with a split ratio of 1:3 at an injector temperature of 200 °C. The oven temperature was isothermally set to 40°C for 6 min; the interface, converting CH_4_ into CO_2_, was running at 940°C. δ^13^C values were referenced against a CO_2_ gas (Air Liquide) and the analytical precision was assessed by repeated measurements of a CH_4_ gas standard (Air Product), both with known δ^13^C value. Each sample was analyzed in duplicates, which revealed an overall precision of <0.5‰. The carbon isotope composition is expressed relative to the VPDB standard:

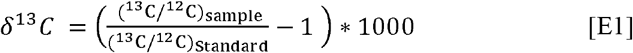

### DIC concentration and stable carbon isotope analysis

To analyze concentration and δ^13^C values of DIC, 550 µL of the incubation medium was filtered (Minisart, 0.22 µm, High Flow PES) into 12-mL Exetainer® vials (Labco), which prior were filled with 100 µL 45% phosphoric acid and purged with helium gas. and After equilibration overnight, the released headspace CO_2_ was analyzed using a DeltaRay IRIS with URI connect and a Cetac ASX-7100 Autosampler (Thermo Fisher Scientific). The δ^13^C values were calibrated against an isotopically known CO_2_ reference gas and the precision (<0.5‰) was assessed by regular measurements of isotopically known sodium bicarbonate (Sigma-Aldrich). The resulting CO_2_ volume was converted into a DIC concentration via the gas constant at 25°C and corrected by the instrument’s recovery rate of 86%. The carbon isotope composition is expressed as above relative to the VPDB standard (E1).

### Sulfide and sulfate concentrations

For the analysis of sulfur species 1 mL aliquot of filtered incubation medium was fixed with 0.5 mL 100 mM zinc acetate until measurements. For sulfide measurements, the formed zinc acetate was homogenized in the solution. In subsamples, sulfide was measured using the Cline assay (Cline, 1960) and photometric measurement at 670 nm after a 30-minute reaction period in the dark. For sulfate measurements, fixed samples were diluted 1:50, and sulfate concentrations were measured by ion chromatography (930 compact IC flex, Metrohm with Metrosep A Supp 4 150/4.0 column).

### Lipid extraction and derivatization

After incubation, sediment slurries were centrifuged, freeze-dried (0.5-1 gram dry weight, g_dw_), and extracted via a modified Bligh and Dyer protocol (Sturt et al., 2004). Prior to extraction, behenic acid methyl ester and 1-nonadecanol were added as internal standards. From the total lipid extract (TLE), fractions of fatty acids (FAs) and neutral lipids (NLs) were obtained by saponification with 6% KOH (Sigma-Aldrich) in methanol (Sigma-Aldrich) reacting for 3h at 80°C and subsequently extracted three times with *n-*hexane under basic and acidic conditions, respectively. Both fractions were dried under a stream of nitrogen and stored at −20°C until further workup. FAs were derivatized with 10% BF_3_ in MeOH (VWR) at 70°C for 1hforming fatty acid methyl esters (FAMEs) and extracted three times before they were dried under a stream nitrogen and stored at −20°C before measurements within the next 5 days. NLs, containing hydrocarbons and alcohols, were derivatized using BSTFA (Sigma-Aldrich) in pyridine (anhydrous, 99.8%, Sigma-Aldrich) for 1h at 70°C forming TMS derivatives of the alcohols. TMS derivatives were measured upon the same day to avoid hydrolysis.

### Lipid analysis via GC-FID, GC-MS and GC-IRMS

Lipids were quantified relative to the internal standards using gas chromatography coupled to a flame ionization detector (GC-FID; Trace GC Ultra, Thermo Scientific). Before, lipids were qualitatively confirmed by coupled GC-mass spectrometry (GC-MS; Trace 1310 + ISQ 700, Thermo Scientific) equipped with an EI source set at 70 eV. The single quadrupole acquired masses in EI full scan from m/z 50-850. δ^13^C values were determined in duplicate using GC-IRMS (Trace GC Ultra coupled to a GC-IsoLink and connected via a ConFlow IV interface to a Delta V Plus IRMS, Thermo Scientific). All GCs were set to splitless mode injection (1 min) at 310°C, were equipped with the same Restek Rxi-5□ms column (30□m□×□250□μm□×□0.25□μm, Restek) and used the same temperature program, which was: initial oven temperature held at 60°C for 1□min, increase to 150°C at a rate of 10°C min^−1^, raise to 310°C at a rate of 4°C min^−1^ and final held at 310°C for 30□min for NLs and 20 min for FAMEs. The carrier gas for all GC measurements was helium with a constant flow rate of 1.0□mL□min^−1^, except the GC-IRMS where 1.2 mL min^−1^ were applied. For GC-IRMS measurements, compounds were transformed into CO_2_ in a combustion reactor at 940°C after separation. Calibration of the instrument was performed against an isotopically known CO_2_ reference gas and the precision was assessed by regular measurements of a C_20_–C_40_ *n-*alkane standard, which has a long-term variance of ±0.5‰. δ^13^C values of each fraction from duplicate measurements showed deviations of <1‰. δ^13^C values of FAMEs and TMS-derivatives were corrected for additional carbon introduced during derivatization and are reported in delta notation (δ^13^C) relative to VPDB, E1). Representative visualization of GC-FID runs of the NL and FA fraction are shown in the supplementary material (Figure S1).

### Rate determination of SO_4_^2-^ consumption, H_2_S production, DIC consumption and CH_4_ oxidation

SO_4_^2-^, H_2_S and DIC consumption or production rates (Table 1) were calculated by dividing the measured change in concentration in mM (Table 1) by the volume of the liquid (0.1 L) in each incubation bottle, the time of sampling and its corresponding dry weight to yield a rate in µmol g_dw_^−1^ day^−1^.

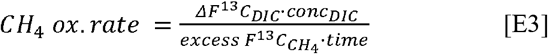

Methane oxidation rates were calculated based on the change in δ^13^C values of DIC in the DIC with ^13^CH_4_ experiment, where the initial δ^13^CH_4_ value at day 0 was +10,000 ‰. During the 30-day incubation period, there was a detectable increase in the concentration and δ^13^C value of DIC, leading to final values of +9.1 mM and +1,140 ‰, respectively (Figure S2, Table S2). The enrichment in ^13^C was used to calculate the corresponding CH_4_ oxidation rate using the equation [E3] (Kellermann et al., 2012):

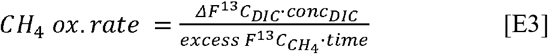

In the equation, δF^13^C represents the change in the fraction of ^13^C, while the excess F^13^C_CH4_ represents the initial fraction of ^13^C. Methane oxidation rates after 10, 20 and 30 days can be found in Table 1.

### Determination of assimilation rates and lipid turnover times

The assimilation rate of IC (assim_IC_; Kellermann et al., 2012; Wegener et al., 2012) was calculated by multiplying average lipid concentrations (conc_lipid_) with the increase of ^13^C in the lipids (δF^13^C_lipid_) relative to the non-labeled samples, and divided by the fraction of ^13^C in the incubation medium (F^13^C_medium_) and the incubation time (t), as shown in the following equation:

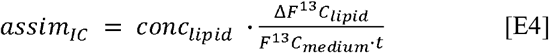

In this equation, F^13^C refers to the proportion of ^13^C in both the lipid and the medium, calculated from the respective stable carbon isotopic ratios 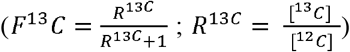. For DI^13^C with CH_4_, we used the measured δ^13^C_DIC_ values, assuming neglectable change (< 1%) in the δ^13^C_DIC_ pool over the course of the incubation. In the experiments of DIC with ^13^CH_4_, we used the average δ^13^C_DIC_ value between day 0 and each day of sampling (10, 20 or 30), because in this treatment δ^13^C_DIC_ increases significantly over time and the discrete measurements at each day of sampling do not reflect the average δ^13^C_DIC_ value.

For calculation of lipid turnover times, we divided the F^13^C_medium_ value by the overall difference in δF^13^C_lipid_ and multiplied with the time (t).

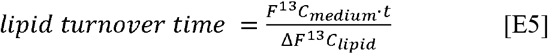

### DNA extraction and short-read sequencing

DNA was extracted from pellets of 250 mg of sediment samples using the DNeasy PowerSoil Pro kit (Qiagen). Total DNA yield per sample, determined by fluorometric DNA concentration measurement, ranged from 226.5 µg and 465 µg (4.53 and 9.3 ng/µl; 50 µl per sample). Samples were sequenced at the LGC Genomics GmbH (Berlin, Germany). Samples were sequenced as 2□×□150 bp paired-end reads on an Illumina MiSeq sequencing platform. Between 19,152,757 (Start, day 0) and 20,850,907 (End, day 30) raw reads were obtained.

### Analysis of community composition

Metagenomic raw reads were first quality checked using FastQC v0.11.4 and MultiQC v1.13 (https://github.com/s-andrews/FastQC; https://github.com/MultiQC/MultiQC) with default settings. Low quality reads and adapters were removed from the dataset using BBDuk of the BBtools package v38.98 (https://sourceforge.net/projects/bbmap/; parameters: minlength=50 qtrim=rtrimq=20). Microbial community composition based on 16S rRNA gene abundance was calculated with phyloFlash v. 3.3b1 (https://hrgv.github.io/phyloFlash/).

### Tetra-labeled oligonucleotides fluorescence in situ hybridization (Tetra-FISH)

At the start and the end of the incubation (day 0 and 30) 3 mL of sediment slurries were fixed for 2 hours using formaldehyde (2.5%) (Sigma-Aldrich) at room temperature. After fixation, the samples were washed with phosphate-buffered saline (PBS) at pH 7.2 and stored in a 1:1 mixture of PBS and ethanol at −20°C until further analysis. Tetra-FISH probes were used to visualize the samples, modified the previously established protocol (Pernthaler et al., 2002, 2001). A formamide concentration of 50% was used for probe ANME2-538 with the sequence (5’ to 3’) GGCTACCACTCGGGCCGC targeting the ANME-2 cluster with the fluorochromes Alexa Fluor 594 (Treude et al., 2005). Formaldehyde-fixed sediment samples stored in PBS-ethanol were diluted ten time with PBS-ethanol and sonicated on ice using a type MS72 probe (Sonopuls HD70; Bandelin, Berlin, Germany; 8 cycles for 30 s with 86% amplitude). 100 µL of the mixture were filtered onto 0.2□µm pore size polycarbonate filters (GTTP, Millipore, Eschborn, Germany) (Ravenschlag et al., 2000). Filters were cut into 10 sections, and each filter section was embedded in 0.1% (w/v) agarose, and dried face down onto a parafilm-covered plastic plate at 37 °C for 30 mins. Microbial cell walls were permeabilized with lysozyme solution (10 mg mL^−1^ in 0.05 M EDTA, 0.1 M Tris-HCl (pH 7.5, Fluka). Filter sections were incubated at 37°C for at 60 min followed by incubation with proteinase K solution (0.45 mU mL^−1^ proteinase K in 0.05 M EDTA, 0.1 M Tris-HCl (pH 7.5), 0.5 M NaCl, Fluka) at room temperature for 5 min. Hybridization was performed for 3 h at 46°C in hybridization chambers using a mixture of 5 µL probe working solution (8.4 pmol µL^−1^) and 45 μL of hybridization buffer (900 mM NaCl, 20 mM Tris-HCl (pH 8.0), 0.01% wt. vol^−1^) sodium dodecyl sulfate (SDS), and 50% (vol/vol) formamide) for each filter section. Following the incubation, the filter sections were washed in 50 ml of pre-warmed washing buffer (28mM NaCl, 5 mM EDTA (pH 8.0), 20 mM Tris-HCl (pH 7.5), and 0.01% (wt. vol^−1^) SDS) at 48°C for 15 min. Filter sections were subsequently washed in ultrapure water (MQ; Millipore) and 96% (vol/vol) ethanol, and air-dried on blotting paper (Whatman, UK). Dried filter sections were mounted with mixtures of CitiFluorAF1 (CitiFluor Ltd., London, UK) and Vectashield (Vector Laboratories, Burlingame, CA, USA) containing 1 µg mL^−1^ DAPI (4′, 6-diamidino-2-phenylindole; Sigma-Aldrich, Steinheim, Germany).

### AI usage, data analysis and data visualization

We employed large language models (LLMs), specifically OpenAI’s ChatGPT, for primary code development with respect to data evaluation and or creation of figures. For the quantification of lipids, python packages “pyGC_FID_processing” (Zander, 2024) were used. To validate compound quantification, we manually integrated a subset of chromatograms to compare them with the automated approach. Data analysis generally was carried out with R Studio (RStudio Team, 2024) and Tidyverse (Wickham et al., 2019) where figures were created too. Figure design was finalized with the vector graphic editor Inkscape V1.4 (86a8ad7, 2024-10-11).

## Results

In this study, we used SIP experiments with ^13^CH_4_ and DI^13^C to distinguish between the assimilation of methane or IC into lipid biomass of AOM community members in active cold seep sediments of the Astoria Canyon, Oregon.

### Analysis and visualization of the natural microbial communities

To characterize the microbial community of the cold seep sediment, we extracted and sequenced the total DNA metagenome, and taxonomically analyzed the recovered 16S rRNA gene sequences (Figure 1, A). The relative abundance of the different microbial groups remained largely stable throughout the incubation, with changes of less than 3% on the phylum and order levels, and less than 1.5% on family level (Table S1). Based on the 16S rRNA gene reads, *Halobacteriota* archaea accounted for 22% of the total community at the phylum level, with *Methanosarcinales* representing 21% at the order level. Within this order, ANME-2c was the most abundant, accounting for 10%, followed by ANME-2a/b with 6% (Figure 1 A), Table S2). Desulfobacterota constituted 15% at the phylum level, with *Desulfobacterales* and *Desulfobulbales* representing 8% and 6% of the total microbial community. Within these orders, *Desulfosarcinaceae* was the most abundant family, accounting for 6% of the total microbial community (Figure 1A, Table S1). Other major phyla included Campylobacterota (9%), Proteobacteria (6%), Bacteroidota (6%) and Chloroflexi and Actinobacteriota (5% each) (Figure 1A, Table S1).

**Figure 1:**
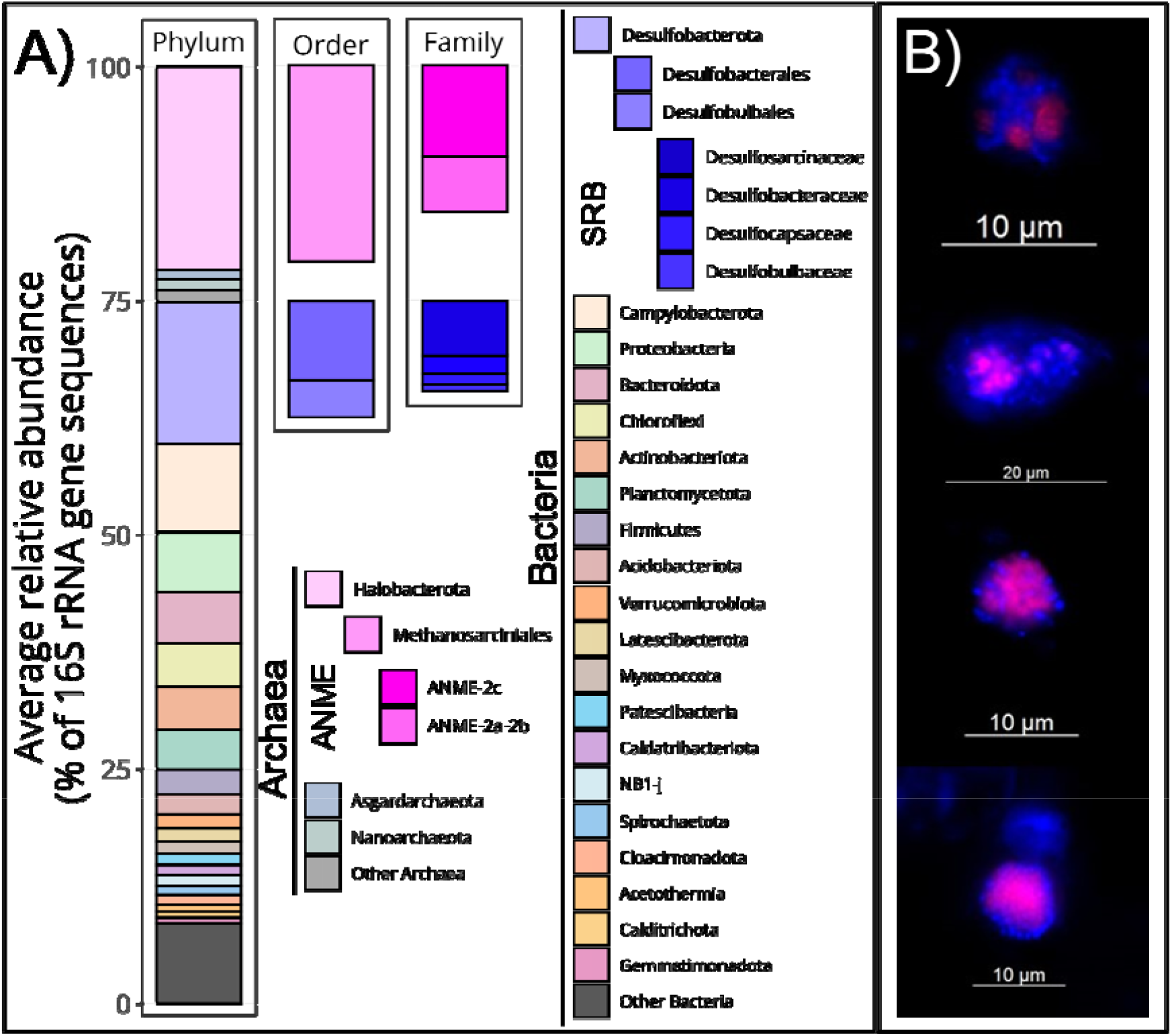
Community analysis and AOM aggregates vizualiation(A) Communtiy composition of the cold seep sediment used in the SIP experiment. Avarage relative abundances of 16S rRNA gene sequences obtained from the metagenome at the start and the end of the SIP experiments are shown (n=2). The panels (from left to right) display annotations at the phylum level of all classified sequences, followed by sequences classified as ANME and SRB at the order and family levels. ANME members are highlighted in magenta, and SRB known to form consortia are highlighted in blue. (B) Fluorescence micrographs (Tetra-FISH) of the cold seep sediments. Images show cells hybridized with the xxx probe specific to ANME-2 (magenta), sourrounded by SRB cells only stained with DAPI, together forming the AOM consortia in various shapes and sizes.

We performed Tetra-FISH analysis at the start and end of the incubation to visualize the ANME-2/SRB consortia (Figure 1 B), Figure S1). A specific ANME-2 probe stained archaeal DNA in magenta, while all DNA was stained with DAPI (blue). Shape and distribution of ANMEs and SRBs in the consortium generally correspond to previously described ANME-2/SRB consortia observed with FISH microscopy (Knittel and Boetius, 2010; Metcalfe et al., 2021). The relatively small size of the observed consortia likely resulted from the harsh ultrasonication step during sample preparation, which fragmented larger aggregates. Nonetheless, the observation of AOM-SR consortia supports the presence and activity of ANME-2 and their SRB partners.

### Lipid composition and δ_13_C values of the original cold seep sediment

Lipid concentrations and their δ^13^C values of the original sediment used for the SIP experiment are shown in Figure 2 and Table S3. The sediment is rich in crocetane (1.3 µg g_dw_^−1^; δ^13^C −100‰), archaeol (Ar, 4.0 µg g_dw_^−1^, −91‰), sn2-hydroxy-archaeol (sn2-OH-Ar,7.4 µg g_dw_^−1^, −101‰) and sn2-phytanyl-mono-alkyl-glycerol ether (sn2-phy MAGE, 1.2 µg g_dw_^−1^, −99‰), which all show strong depletion in δ^13^C. The resulting ratio of sn2-OH-Ar to archaeol is 1.9. The following lipids produced by the partner SRB are also highly abundant but have less negative δ^13^C values: MAGE C_16:1ω5c_ (1.3 µg g_dw_^−1^, −66 ‰), dialkyl glycerol ether (DAGE) C_32:2a_ (5.6 µg g_dw_^−1^, −77 ‰), and FAs like C_16:1ω5c_ (20.7 µg g_dw_^−1^, −61 ‰) and cyC_17:0ω5,6_ (1.1 µg g_dw_^−1^, −59 ‰) (Figure 2, Table S3). Other abundant bacterial lipids have more positive δ^13^C values: C_14:0_ (9.9 µg g_dw_^−1^, −33‰), iC_15:0_ (7.3 µg g_dw_^−1^, −38‰), aiC_15:0_ (5.3 µg g_dw_^−1^, −38‰), C_16:0_ (23.4 µg g_dw_^−1^, −33‰), and C_18:1ω7c_ (7.0 µg g_dw_^−1^, −32‰). Moreover, the sediment shows strong marine input of algal biomarker with typical isotopic compositions such as phytol (10.9 µg g_dw_^−1^, −29‰), cholesterol (5.3 µg g_dw_^−1^, −26‰), sitosterol (7.0 µg g_dw_^−1^, −26‰) and dinosterol (4.0 µg g_dw_^−1^, −24‰).

**Figure 2:**
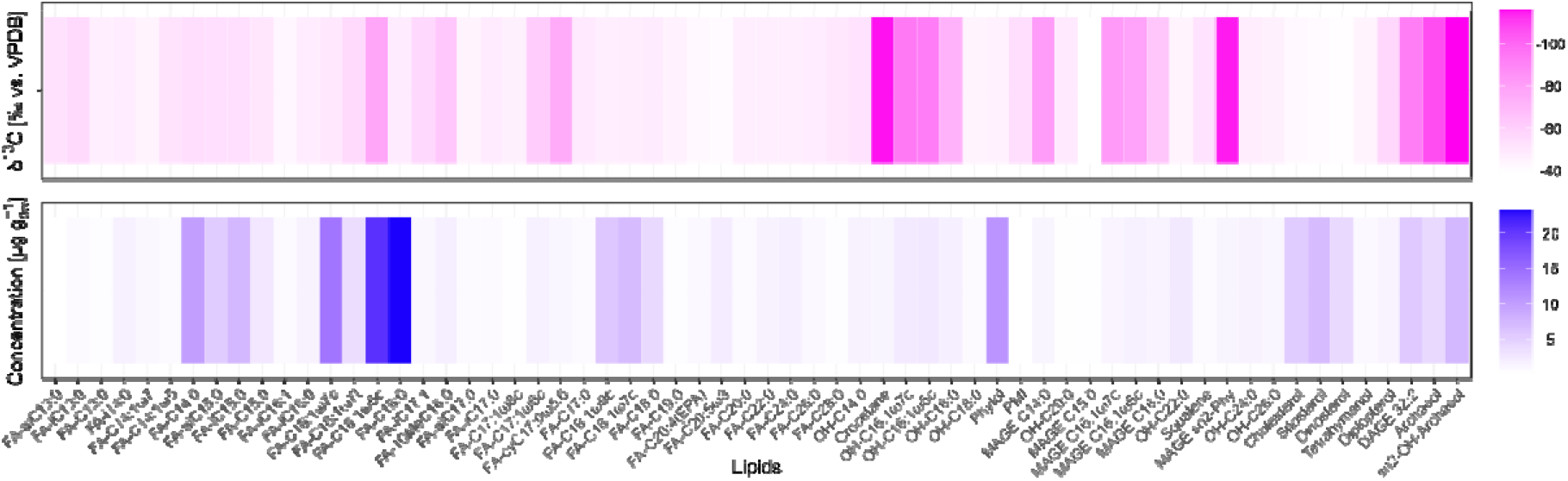
Heat map showing the concentrations (magenta; in µg g_gw_^−1^) and δ_13_C values (blue; in ‰ vs. VPDB) of fatty acids, alcohols and hydrocarbons extracted from the incubated sediment. FA: fatty acid, MAGE: monoalkyl glycerol ether. DAGE: dialkyl glycerol ether. OH: Hydroxy, cy: cyclic, i: iso, ai: anteiso, C: carbon chain length.

### Development of AOM activity during incubation

The geochemical data (δ^13^C values of CH_4_ and DIC, concentrations of DIC, SO_4_^2-^ and H_2_S) from the SIP experiment show the imprint of sulfate-dependent AOM as the dominant microbial process in our experiments (Table 1, Figure 3). Because methane was provided in excess (1 bar) its concentration was not measured. In the ^13^CH_4_-labeling experiment, the δ^13^C-DIC values increased by 1140‰ within 30 days, giving a calculated methane oxidation rate of approximately 4.4 µmol g_dw_□^1^ d□^1^. In the same experiments, sulfate decreased by 4.8±0.3 µmol g_dw_□^1^ d□^1^ (mean and sd, n=3), and sulfide increased up to 5.1±0.7 µmol g_dw_□^1^ d□^1^ (mean and sd, n=3). The slightly higher sulfate reduction than methane oxidation rates can be explained by additional organoclastic sulfate reduction which was 0.x … In all treatments the increase in DIC concentrations was higher than the methane oxidation rates (8.1±1.1 µmol g_dw_□^1^ d□^1^) suggesting dissolution of inorganic carbon. Notably, in the DI^13^C treatments, the δ^13^C-CH_4_ values increased by 60‰. Such label transfer into the CH_4_ pool is known due to the reversibility of the intracellular reactions during AOM (Holler et al., 2011).

**Figure 3:**
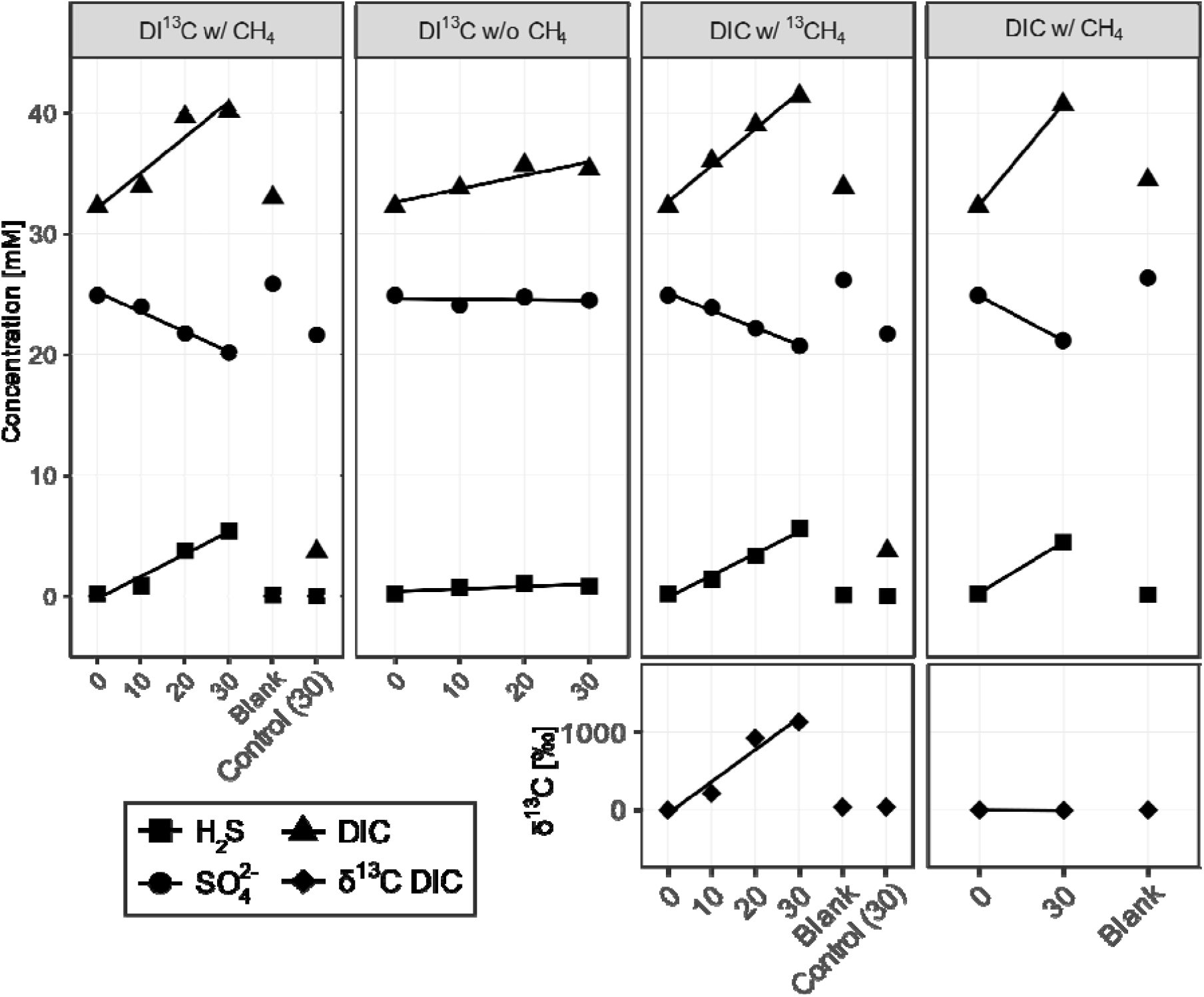
Development of chemical parameters in the four main treatments between 0 and 30 days. First upper four panels show dissolved inorganic carbon (DIC), sulfate (SO_4_^2-^) and hydrogen sulfide (H_2_S) concentrations in mM, the two lower t panels display δ_13_C values of DIC in ‰. All treatments with (w/) methane (CH_4_) show an increase over time, while the treatment without (w/o) CH_4_ does not. The treatment with ^13^C-labeled CH_4_ shows steady label transfer into the DIC pool.

### Lipid-specific labeling with DI^13^C and ^13^CH_4_

Throughout the incubation, concentrations of individual lipids remained largely stable, which is consistent with the slow growth of the AOM consortia. Nevertheless, diagnostic lipid biomarkers of ANME-2 and SRB showed substantial assimilation of the supplied ^13^C-labeled carbon sources (Figure 4, Table S4). We observed the most pronounced 13C assimilation in the SRB specific fatty acids C_16:1ω5c_ and cyC_17:0ω5,6_ with changes in the stable carbon isotopic composition (Δδ^13^C-values) of +393‰ and +258‰, respectively, in the DI^13^C w/CH_4_ experiment after 30 days. Other fatty acids (C_14:0_, C_16:1_ ω_7c_ and C_16:0_) were also strongly ^13^C-labeled reaching Δδ values of +172‰, +197‰, and +129‰, respectively. Of the archaeal lipids, Δδ^13^C-values were highest for crocetane (+126‰) and MAGE sn2-phy (+78.4‰). In experiments without methane (DI^13^C w/o CH_4_), the DI^13^C-label assimilation was minimal: archaeal lipids showed Δδ^13^C-values between +0.3‰ and +10.4‰, while bacterial lipids’ ranged from +6.6‰ to +13.5‰. In experiments with ^13^CH_4_ addition, SRB-specific lipids reached Δδ^13^C values of +55.6‰ (C_16:1_ ω_5c_) and +32.8‰ (cyC_17:0 ω5,6_) and archaeal lipids reached Δδ^13^C values of +36.9‰ for crocetane and +27.7‰ for archaeol. This suggests DIC as primary carbon source of both, ANME and SRB, in DI^13^C and ^13^CH_4_ treatments because the ^13^C-labeling strength of methane was much higher than that of DIC (δ^13^C-CH_4_ ∼10.000‰; δ^13^C-DIC ∼5000‰, Table 1). Bacterial fatty acids rather produced by ancillary bacteria such as *ai*C_15:0_ and *i*C_15:0_ showed Δδ^13^C values of +21.3 and +14.1‰, respectively, in the DI^13^C w/o CH_4_ treatment. These values were only slightly higher in the DI^13^C with CH_4_ experiment (Δδ^13^C values +37.8 and +28.8‰ respectively).

**Figure 4:**
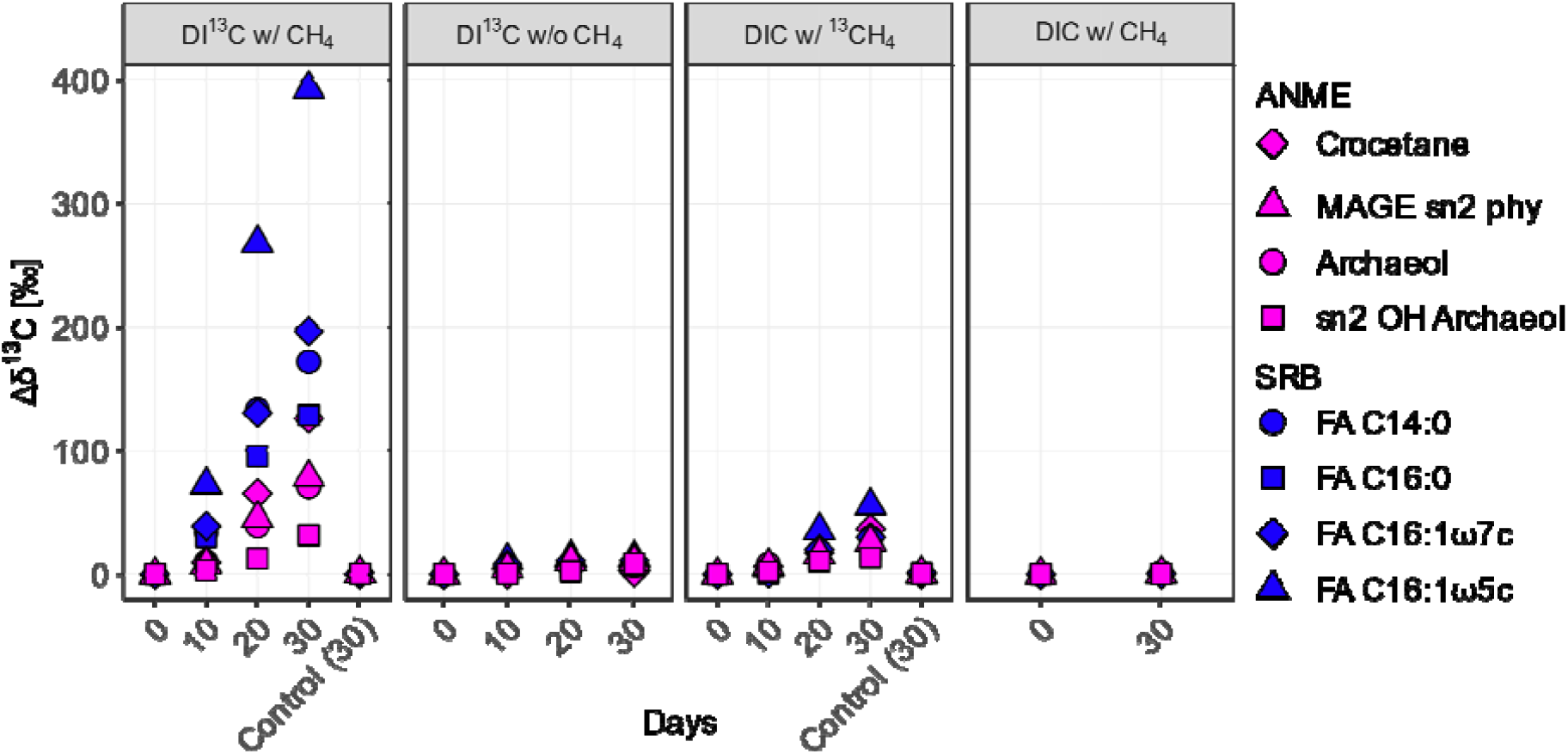
Shifts in δ_13_C values (Δδ_13_C) of lipid biomarkers in the SIP incubation experiments. Each panel represents one treatment over time, from left to right: DIC with ^13^CH_4_, DI^13^C with CH_4_, DI^13^C without CH_4_ and non-labeled DIC with CH_4_. Colors represent lipids diagnostic for ANME (magenta) and the most strongly ^13^C-labeled lipids diagnostic for SRB (blue) grouped by compound classes, i.e., FAs, MAGEs and DAGEs. Shapes represent individual lipid biomarkers and are consistent within each microbial group. FA: fatty acid. MAGE: monoalkyl glycerol ether. DAGE: dialkyl glycerol ether. OH: Hydroxy.

### DIC and methane assimilation rates into AOM specific lipids

We calculated assim_IC_ for treatments with labelled DI^13^C and all time points according to the equation E4. In the DI^13^C w/CH_4_ incubations, SRB lipids showed the highest assim_IC_ values. For example, C_16:1_ω_5c_ reached 13.6 µg C g_dw_^−1^ y^−1^ by day 30 (Figure 5, Table S5) and C_16:0_ reached 5.3 µg C g_dw_^−1^ y^−1^. Archaeal lipids also assimilated substantial ^13^C-label in the same DI^13^C w/CH_4_ treatment by day 30: archaeol reached 0.49 µg C g_dw_^−1^ y^−1^ and sn2-OH-archaeol 0.24 µg C g_dw_^−1^ y^−1^. In the treatment without methane (DI^13^C w/o CH_4_, Figure 5, Table S5), ^13^C–DIC assimilation into lipids dropped substantially. After 30 days, assim_IC_ in SRB lipids was by average 94□±□4% (mean and SD, n=4) lower, and in ANME-2 archaeal lipids 87□±□11% (mean and SD, n=4) lower, compared to the conditions with methane. By comparison, the heterotrophic background community (e.g., bacteria marked by fatty acids *ai*C_15:0_, *i*C_15:0_, and C_18:1ω7c_) showed overall lower assim_IC_ values (0.2–0.4 µg C g_dw_^−1^ y^−1^). Notably, ^13^C-assimilation into these lipids was less affected by the absence of methane (only ∼57□±□15% (mean and SD, n=3) reduction without methane).

**Figure 5:**
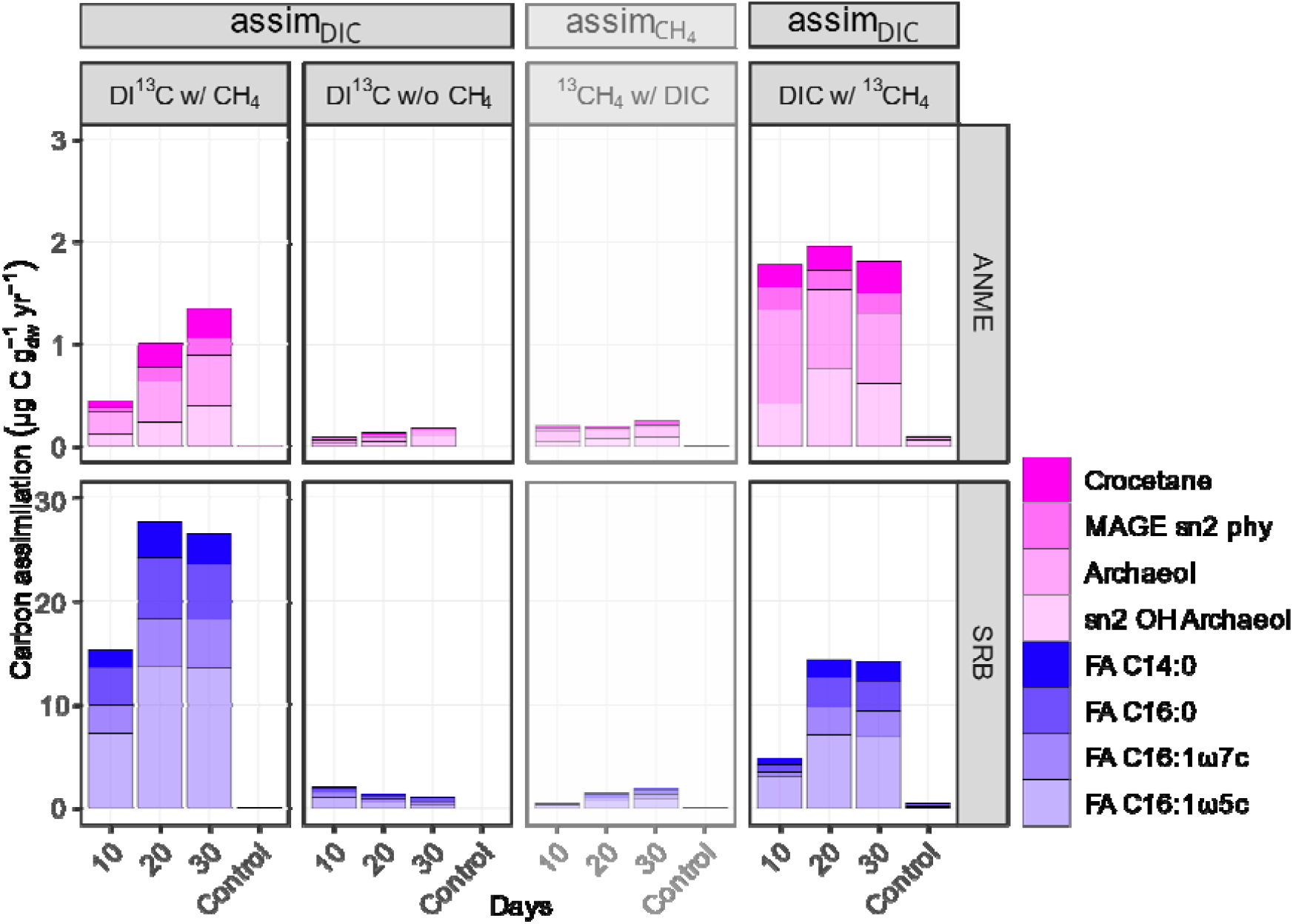
Incorporation of carbon from dissolved inorganic carbon (DIC) and methane into diagnostic lipids of AOM-associated microorganisms across stable isotope probing (SIP) treatments. Vertical panels represent individual SIP treatments: from left to right (1) DI^13^C with CH_4_, (2) DI^13^C without CH□. Both treatments show inorganic carbon assimilation (assim_IC_) rates. (3) ^13^CH□ with DIC. This treatment shows potential methane carbon assimilation (assim_CH4_) with reduced opacity. (4) DIC with ^13^CH_4_. This treatment shows inorganic carbon assimilation (assim_IC_). Horizontal panels separate microbial groups: anaerobic methanotrophic archaea (ANME, top) and sulfate-reducing bacteria (SRB, bottom). Bar heights indicate carbon assimilation in µg C g_dw_ yr^−1^ into diagnostic lipids after 10, 20, and 30 days. Where available, 30-day values for killed controls are shown for comparison. Lipid classes include fatty acids (FA), monoalkyl glycerol ethers (MAGE), dialkyl glycerol ethers (DAGE), and hydroxy (OH) groups. The shading represents individual compounds.

To assess the role of methane as a carbon source, we calculated assim_CH_□ values exclusively for treatments where CH□ was ^13^C-labeled (Figure 5, lesser opacity, Table S6). Such calculations were not considered in any lipid SIP study yet and values for SRB are only hypothetical due to their sole autotrophic carbon fixation lifestyle. These calculations revealed that carbon assimilation from methane would be substantially lower than from DIC, particularly in archaeal lipids. For instance, in the DIC w/^13^CH_4_ treatment, only 0.1 µg gdw^−1^ y^−1^ of archaeol could be attributed to methane as a carbon source, which is similar to the assim_CH_□ value for archaeol in the killed control of the same treatment after 30 days. To compare assim_IC_ and assim_CH_□ rates during the treatment DIC w/^13^CH_4_, we used the averaged δ^13^C-DIC values which increased to 1129‰ during the 30 days of experiments to calculate assim_IC_. Those derived assim_IC_ values were on the same magnitude than those during the DI^13^C w/CH_4_ treatments (Figure 5, right panel, Table S5). Values for ANME markers crocetane, sn2-phy MAGE, archaeol and sn2-OH-Ar were between 0.2 and 0.7 µg C g_dw_^−1^ y^−1^ and values for SRB FA C_16:1 ω5c_ and C_16:0_ were 2.9 and 6.9 µg C_dw_ g ^−1^ y^−1^ (Table S5).

Temporal development of individual and summed assim_IC_ values is different for ANMEs compared to SRBs (Figure 5, Table S5). In the DIC w/^13^CH_4_ experiment, the assim_IC_ into summed ANME lipids was already substantial by day 10 and remained stable above 1.5 µg gdw^−1^ y^−1^ until day 30. By contrast, in the DI^13^C w/CH_4_ experiment, ^13^C assimilation into ANME lipids started lower and steadily increased between day 10 and day 30 to close to 1.5 µg gdw^−1^ y^−1^. SRB specific showed similar developments of assim_IC_ values during the course of both experimental types. In the treatment DI^13^C w/CH_4_, summed bacterial FAs showed values of around 15 µg gdw^−1^ y^−1^ after 10 days which increases to ∼30 µg gdw^−1^ y^−1^ after 20 days. After 30 days, their assim_IC_ remained largely stable. The same temporal pattern of the assim_IC_ values was observed for the DIC w/^13^CH_4_ treatment for the partner bacteria specific lipids with the exception of assim_IC_ values being half as high.

### Lipid turnover times during SIP with DI^13^C

ANME lipid turnover times obtained according to E5 ranged from 3.6 to 14.5 years, while those for the lipids of their SRB partner were shorter, around 1 to 7 years (Table S7). At the final incubation point, archaeal lipids crocetane, archaeol, and sn2-Phy-MAGE had turnover times of 3.6, 5.9 and 6.5 years. In contrast, sn-2-OH-Ar exhibited a notably longer turnover time of ∼13.3 years. Among the SRB lipid markers, the fatty acid C_16:1ω5c_ displayed the shortest turnover time (1.2 years). Other SRB lipid markers showed intermediate turnover times (∼1.8 – 7.8 years), including MAGE C_16:1ω5c_, DAGE C_32:2_, and the more ubiquitous fatty acids C_14:0_ and C_16:0_. These longer turnover times might indicate either a higher fossil lipid contribution or, particularly in the case of C_14:0_ and C_16:0_, reduced biomarker specificity since various other bacterial groups also synthesize these fatty acids.

## Discussion

Metagenomic and microscopy analyses provide evidence that the microbiome of the methane-rich cold seep sediment in Astoria Canyon is dominated by anaerobic methanotrophic archaea of the ANME-2 clade and their syntrophic SRB partners (Figure 1). Together, ANME-2c and ANME-2a/b comprised 15.6% of the community at the family level, while *Desulfobacterota* level - particularly *Desulfosarcinaceae* - accounted for 15.1% on phylum level (Table S2). The co-existence of ANME-2a/b and ANME-2c with SRB at an approximate 1:1 level is often observed at methane seep locations with high AOM activity like the Eel River Basin (Pernthaler et al., 2008), Hydrate Ridge (Boetius et al., 2000; Treude et al., 2003), and the Black Sea (Blumenberg et al., 2004). The shape and spatial arrangement of AOM consortia in those environments closely resembles the Tetra-FISH micrographs of the Astoria Canyon sediment community presented here (Figure 1). Strongly ^13^C-depleted archaeol, sn2-OH-archaeol, and crocetane as diagnostic lipid biomarkers further supports the dominance of ANME-2 (e.g. Blumenberg et al., 2004; Elvert et al., 2005; Niemann and Elvert, 2008). Additionally, a sn2-OH-Ar to archaeol ratio of 1.9 in our samples is consistent with ANME-2 and distinct from ANME-1SRB specific lipids are also abundant, i.e. C_16:1ω5c_, cyC_17:0ω5,6_ and their respective MAGE, alcohol and DAGE derivates of C_16:1ω5c_, and have low δ^13^C-values, although not as strongly ^13^C-depleted as the lipids produced by ANME. This pattern is consistent with previous findings in methane-rich seep sediments (e.g. Hinrichs et al., 2000; Elvert et al., 2003, 2005; Blumenberg et al., 2005), which authors also found a clear isotopic difference between more ^13^C-depleted ANME lipids and slightly less ^13^C-depleted SRB lipids. The heterotrophic background community, including members of *Campylobacterota* and *Bacteroidota* closely resembles those found in other seep ecosystems (Pop Ristova et al., 2015; Zhu et al., 2022). These organisms produce C_18:1ω9c_, *ai*C_15:0_ and *i*C_15:0_, which display moderately low carbon isotope values (δ^13^C = −30 to −38‰) and based on our SIP experiments with DI^13^C are largely formed by methane-independent DIC assimilation (Table S5). This observation suggests that the heterotrophic community relies on a mix of degraded sedimentary organic matter and necromass of the AOM core community (Zhu et al., 2022).

Despite methane being ^13^C-labeled more strongly (δ^13^C ∼10,000‰) than DIC (δ^13^C ∼5,000‰), we found that assim_IC_ was about one magnitude greater than ^13^CH_4_ assimilation (assim_CH4_) into diagnostic lipids of both ANME and SRB (Figure 5, Table S5 and S6), The minor assim_CH4_ values observed likely reflect assimilation of methane-derived DIC, as DI^13^C assimilation calculated for the DIC w/^13^CH_4_ treatment were comparable to those in the DI^13^C w/CH_4_ treatment (Figure 5, Table S5). These results underline previous SIP studies using only ^13^C-labeled methane (Bertram et al., 2013; Blumenberg et al., 2005; Jagersma et al., 2009), and refine the conclusions of Wegener et al. (2008), whose long-term incubations may have masked distinct assimilation pathways due to equilibration between methane- and CO_2_ pools. The ANME-2/SRB consortia therefore mainly use IC as carbon substrate for their lipid biomass. The assimilation of IC into lipids attributed to SRB were approximately 8 times higher after 30 days than those attributed to ANME archaea (Figure 5, Table S5, (Figure 5, Table S5, summed assim_IC_ values for archaeal vs. bacterial lipids (n=4)), resulting in shorter turnover times for SRB (1–7.8 years) compared to ANME lipids (3.6–13.3 years). This is counterintuitive, as SRB and ANME typically occur in AOM consortia in a ratio around 1:1 which suggests similar growth rates (Scheller et al., 2016). A likely explanation for the underestimated ANME lipid production is that the ANME continuously dilute the ^13^C-DIC signal though the intercellular production of methane-derived CO_2_. In other words, during the DI^13^C labeling experiment, ANME cells experience the externally measured average DI^13^C value, but are internally exposed to a ^13^C-diluted CO_2_ pool due to continuous AOM. To quantify this effect, we recalculated assim_IC_ using the δ^13^C-DIC values from the ^13^CH_4_ experiment, where internally methane-derived CO□ dominates. Using this approach, archaeal lipid production rates increase by approximately 1.8 times after 30 days (Figure 5, Table S5, summed assim_IC_ values for archaeal lipids (n=4)), suggesting that the initial estimates likely underestimated their true biosynthetic activity. Nonetheless, even under these assumptions, the calculated production of all archaeal lipids remained far lower than that of bacterial fatty acids (Figure 5). An alternative explanation for the underestimation of archaeal lipid production could be the assimilation of additional but yet undetected carbon sources by the ANME. Previous studies have already investigated whether the addition of small organic compounds such as acetate affects AOM rates and consortia growth, but no positive effects were found (Nauhaus et al., 2002; Wegener et al., 2016a). However, the δ^13^C values of ANME-lipids and biomass are far more negative than those of the background organic matter. If ANME additionally assimilate organic carbon, it would likely need to originate from their syntrophic partner bacteria, possibly in the form of metabolic intermediates. To test this possibility, future studies should target and quantify specific metabolites potentially released by SRB and channeled to the archaeal partner. Identifying such intermediates could significantly refine our understanding of carbon cross-feeding and the metabolic interdependence within AOM consortia.

The temporal development of Δδ^13^C values and carbon assimilation rates in the individual SIP experiments provide insight into lipid biosynthesis pathways and functional roles of these compounds. In the SIP experiment with DI^13^C and methane (Tables S4 and S5), the δ^13^C values ANME-2 lipids MAGE sn2-Phy, archaeol and crocetane, were already strongly enriched after 10 days, and continued to increase throughout the experiment. In contrast, sn2-OH-archaeol, exhibited both, delayed and substantially weaker ^13^C-label incorporation, indicating much longer turnover times. While MAGE sn2-Phy and archaeol are synthesized more directly, the former likely as a by-product of hydrogenation via geranylgeranyl reductase of geranylgeranylglyceryl phosphate (GGGP) prior to the formation of digeranylgeranylglyceryl phosphate (DGGGP) in the archaeol biosynthesis pathway (Caforio and Driessen, 2017; Jain, 2014; Villanueva et al., 2014), the pathway to sn2-OH-archaeol is believed to involve hydration of the 2’,3’ double bond in the sn-2 chain of DGGGP (Mori et al., 2015). The differences in ^13^C-label incorporation in our experiments are line with intermediates and products of the lipid biosynthetic route and hence explain the wide range of lipid turnover times in ANME-2. This also resembles the pattern seen in ANME-1, where the less abundant and highly ^13^C-labeled intermediate phosphatidylglycerol-archaeol is rapidly produced and progressively converted to dominant but less ^13^C-labelled GDGTs (Kellermann et al., 2016). In summary, when interpreting on ^13^C-labeling of lipids, both lipid production and transformation pathways must be considered, especially in slow-growing microorganisms such as AOM consortia.

The immediate and continuous ^13^C-labeling of crocetane is intriguing and points to a fundamental role in ANME-2 cell membranes. To date, no definite biosynthetic pathway has been established for the isoprenoidal hydrocarbon formation in archaea. However, there is growing evidence for squalene biosynthesis via the condensation of two farnesyl pyrophosphate units (Rao and Driessen, 2024; Salvador-Castell et al., 2019), possibly mediated by squalene synthase acquired by lateral gene transfer (Santana-Molina et al., 2020), and followed by hydrogenation to form squalane. By analogy, crocetane may be synthesized through the fusion of two geranyl pyrophosphate units to form fully unsaturated crocetene, which is subsequently reduced (Elvert, 1999). This hypothesis is supported by the detection of intermediate hydrocarbons with one or two double bonds in both natural AOM environments and enrichment cultures (Blumenberg et al., 2004; Elvert et al., 2000; Nauhaus et al., 2007; Pancost et al., 2000). The physiological function of isoprenoidal hydrocarbon in archaea remains under debate. However, current evidence suggest that lipid bilayer-forming archaea insert these non-polar hydrocarbons into the midplane to regulate membrane properties, such as enhancing fluidity and reducing proton permeability (cf. LoRicco et al., 2023; Salvador-Castell et al., 2021a, 2021b, 2020), similar to the roles of steroids in eukaryotes and hopanoids in bacteria. Apolar hydrocarbons may represent a strategy by which bilayer-forming archaea could adapt to extreme environmental and energy constraints without synthesizing bipolar, membrane-spanning, monolayer-forming lipids.

Reports on the biosynthesis of ether lipids, i.e. MAGEs and DAGEs, in SRBs remain scarce. However, using a culture of a mesophilic alkene-degrading sulfate reducer, Grossi et al. (2015) provided compelling evidence for a systematic link between the chain length and the methyl-branching pattern of fatty acids, MAGEs and DAGEs. They proposed an unprecedented biosynthetic pathway via primary fatty acids leading to structurally similar MAGEs and DAGEs in anaerobic bacteria. This appears to extend to SRBs involved in AOM, where fatty acid and MAGE patterns - as well as the side chains of corresponding DAGEs - are systematically congruent (e.g. Elvert et al., 2005; Hinrichs et al., 2000). In our current SIP experiments, the biosynthetic trend is evident through the highest ^13^C-assimilation observed in the fatty acid C_16:1ω5c_, which appears to propagate along the biosynthetic pathway via MAGE finally yielding DAGE C_32:2_ as probable end product (Tables S4 and S5). DAGE C_32:2_ likely consists of two alcohol chains with a C_16:1ω5c_ structure (Elvert et al., 2005), although an incorporation of C_16:1ω7c_ cannot be ruled out, given its abundance and similar ^13^C-depletion in the original sediment (Figures 3 and S1). To date, enzymes involved in the formation of ether lipids in anaerobic bacteria have rarely been characterized. An exception are key enyzmes in *Thermoanaerobacter* (phylum Firmicutes), which can synthesize membrane-spanning lipids by fusion of two *i*C_15:0_ fatty acids and subsequent reduction to form non-isoprenoidal ether lipids (Sahonero-Canavesi et al., 2022). Similar to archaea, the functional role of ether lipid production in bacteria likely relates to enhanced membrane stability. In the context of AOM, this stabilization, especially in cooperation with ANMEs, may contribute to greater energy efficiency.

## Conclusion

We followed the assimilation of ^13^C-label into individual lipids of AOM consortia dominated by ANME-2 and SRB of the order *Desulfobacterales* and *Desulfobulbales* during SIP incubations with DI^13^C or ^13^CH_4_ and calculated lipid-specific assimilation rates and turnover times to identify their carbon source. The data showed that IC is incorporated far more efficiently into lipid biomass than methane carbon in both ANME-2 and their SRB partners. However, SRB lipids exhibited eight times higher carbon assimilation compared to archaeal lipids. Examination of the temporal pattern of ^13^C-label incorporation in experiments with DI^13^C suggests that the low carbon assimilation in archaeal lipids may be due to additional fixation of internally generated CO_2_ from ^13^C-label-free methane before it can be exchanged with the environment. Because this effect still cannot fully account for the observed differences in archaeal versus bacterial lipid production, we propose that ANME-2 rely on another carbon source, for example, small organic compounds provided by their SRB partners. Finally, the step-wise ^13^C-enrichment of ANME- and SRB-derived lipids tracks biosynthetic pathways forming diether lipids in AOM consortia and points towards crocetane as a membrane-modulating isoprenoid hydrocarbon that affects fluidity and proton permeability under low-energy conditions.

## Author Contributions

GW, ME, and LS conceived and designed the study. LS and GW conducted the stable isotope probing experiments. LS performed lipid extraction, compound-specific isotope analyses, and data interpretation. YW carried out DNA extraction, microbial community analysis, and tetra-FISH. YZ contributed to analytical discussions and interpretation. LS, GW, and ME wrote the manuscript. All authors reviewed and approved the final version of the manuscript.

## Acknowledgements

This research was supported by the Deutsche Forschungsgemeinschaft (DFG, German Research Foundation) under Germany’s Excellence Strategy through the Cluster of Excellence ‘The Ocean Floor—Earth’s Uncharted Interface’ (EXC 2077 – 390741603). Sampling during cruise AT50-14 (Seep-DOM) was funded by the U.S. National Science Foundation (NSF, award number 2049517). We thank Jenny Altun, Heidi Taubner, Martina Alisch, and Marco Feo Ziemann for technical assistance. We are also grateful to the captain and crew of the R/V Atlantis and the scientific party of AT50-14 for their invaluable contributions during sampling.

## Data Availability Statement

All raw reads generated in this study have been deposited at ENA - BioProject [number]. Additional data (isotopic values, lipid concentrations) available in the supplementary material and on …

## Ethics Statement

All necessary permissions for field sampling were obtained from the relevant national and institutional authorities. No experiments were conducted on humans or vertebrate animals. Research complied with all applicable international, national, and institutional guidelines for marine scientific research, including those related to the Convention on Biological Diversity and the Nagoya Protocol.

## Supporting Information

Additional supporting information can be found online in the Supporting Information Section

