## Supplemental Material for "Marine cold seep ANME-2/SRB consortia produce their lipid biomass from inorganic carbon"

Supplemental files

**Table S1**: Summary statistics for raw and assembled metagenomes of the unlabeled samples at the start (0 days) and the end (30 days) of the incubation.

| **Sample** | **Unit** | **Start (0 days)** | **End (30 days)** |
| --- | --- | --- | --- |
| **DNA yield** | ng µL^-1^ | 9.3 | 4.5 |
| **Raw reads** | M | 19.2 | 20.9 |
| **Trimmed reads** | M | 18.6 | 20.3 |
| **Total assembly length** | bp | 1,179,793,415 | 1,209,262,363 |
| **L50** |  | 254 | 252 |
| **N50** |  | 369,690,1 | 3,831,154 |
| **Number of contigs** |  | 4,320,921 | 4,471,251 |
| **Max contig length** | bp | 34,093 | 14,664 |

**Table S2**: Summary of 16S rRNA gene sequences recruited from the metagenome at the phylum, order and family levels, used to generate the plot in Figure 2A. The data were obtained from the unlabeled treatment at the start (0 days) and the end (30 days) of the incubation. All values are given as relative abundances (%). AVG indicates the average of the start and the end relative abundance values (n=2).

| **Phylum level** | | | | | |
| --- | --- | --- | --- | --- | --- |
| **Kingdom** | **Phylum** | | **Start** | **End** | **AVG** |
| Archaea | Asgardarchaeota | | 0.9 | 1.3 | 1.1 |
| Archaea | Halobacterota | | 23.1 | 20.3 | 21.7 |
| Archaea | Nanoarchaeota | | 1.1 | 1.1 | 1.1 |
| Archaea | Other Archaea | | 1.4 | 1.3 | 1.4 |
| Bacteria | Acetothermia | | 0.7 | 0.8 | 0.8 |
| Bacteria | Acidobacteriota | | 2.3 | 1.8 | 2.1 |
| Bacteria | Actinobacteriota | | 4.7 | 4.4 | 4.6 |
| Bacteria | Bacteroidota | | 5.3 | 5.7 | 5.5 |
| Bacteria | Caldatribacteriota | | 1.1 | 1.2 | 1.2 |
| Bacteria | Calditrichota | | 0.5 | 0.7 | 0.6 |
| Bacteria | Campylobacterota | | 8.9 | 9.8 | 9.4 |
| Bacteria | Chloroflexi | | 4.7 | 4.6 | 4.6 |
| Bacteria | Cloacimonadota | | 1.0 | 1.0 | 1.0 |
| Bacteria | Desulfobacterota | | 13.7 | 16.5 | 15.1 |
| Bacteria | Firmicutes | | 2.8 | 2.5 | 2.6 |
| Bacteria | Gemmatimonadota | | 0.6 | 0.6 | 0.6 |
| Bacteria | Latescibacterota | | 1.6 | 1.3 | 1.4 |
| Bacteria | Myxococcota | | 1.3 | 1.3 | 1.3 |
| Bacteria | NB1-j | | 1.2 | 1.0 | 1.1 |
| Bacteria | Other Bacteria | | 8.4 | 8.8 | 8.6 |
| Bacteria | Patescibacteria | | 1.0 | 1.3 | 1.2 |
| Bacteria | Planctomycetota | | 4.4 | 4.1 | 4.2 |
| Bacteria | Proteobacteria | | 6.5 | 6.4 | 6.4 |
| Bacteria | Spirochaetota | | 1.0 | 1.0 | 1.0 |
| Bacteria | Verrucomicrobiota | | 1.6 | 1.4 | 1.5 |
| **Order level** | | | | | |
| **Kingdom** | **Phylum** | **Order** | **Start** | **End** | **AVG** |
| Archaea | Halobacterota | Methanosarciniales | 22.3 | 19.5 | 20.9 |
| Archaea | Other Archaea | Other Archaea | 3.0 | 3.4 | 3.2 |
| Archaea | Nanoarchaeota | Woesearchaeales | 1.1 | 1.1 | 1.1 |
| Bacteria | Actinobacteriota | Actinomarinales | 1.7 | 1.3 | 1.5 |
| Bacteria | Chloroflexi | Anaerolineales | 2.8 | 2.7 | 2.8 |
| Bacteria | Bacteroidota | Bacteroidales | 2.4 | 2.8 | 2.6 |
| Bacteria | Calditrichota | Calditrichales | 0.5 | 0.7 | 0.6 |
| Bacteria | Campylobacterota | Campylobacterales | 8.9 | 9.8 | 9.4 |
| Bacteria | Cloacimonadota | Cloacimonadales | 1.0 | 1.0 | 1.0 |
| Bacteria | Desulfobacterota | Desulfatiglandales | 0.9 | 0.9 | 0.9 |
| Bacteria | Desulfobacterota | Desulfobacterales | 7.6 | 9.3 | 8.4 |
| Bacteria | Desulfobacterota | Desulfobulbales | 3.5 | 4.4 | 4.0 |
| Bacteria | Bacteroidota | Flavobacteriales | 2.0 | 1.7 | 1.8 |
| Bacteria | Proteobacteria | Gammaproteobacteria Incertae Sedis | 0.7 | 0.7 | 0.7 |
| Bacteria | Latescibacterota | Latescibacterales | 0.9 | 0.7 | 0.8 |
| Bacteria | Planctomycetota | MSBL9 | 1.5 | 1.7 | 1.6 |
| Bacteria | Actinobacteriota | Microtrichales | 1.2 | 1.0 | 1.1 |
| Bacteria | Other Bacteria | Other Bacteria | 30.2 | 30.6 | 30.4 |
| Bacteria | Firmicutes | Peptostreptococcales-Tissierellales | 1.0 | 0.8 | 0.9 |
| Bacteria | Planctomycetota | Pirellulales | 1.4 | 1.2 | 1.3 |
| Bacteria | Myxococcota | Polyangiales | 1.1 | 1.1 | 1.1 |
| Bacteria | Proteobacteria | Pseudomonadales | 1.1 | 1.0 | 1.0 |
| Bacteria | Proteobacteria | Rhizobiales | 1.0 | 0.8 | 0.9 |
| Bacteria | Spirochaetota | Spirochaetales | 1.0 | 1.0 | 1.0 |
| Bacteria | Acidobacteriota | Thermoanaerobaculales | 1.2 | 0.7 | 0.9 |
| **Family level** | | | | | |
| **Kingdom** | **Phylum** | **Family** | **Start** | **End** | **AVG** |
| Archaea | Halobacterota | ANME-2a-2b | 6.6 | 5.3 | 5.9 |
| Archaea | Halobacterota | ANME-2c | 10.2 | 9.3 | 9.7 |
| Archaea | Halobacterota | Methanosarcinaceae | 1.9 | 1.7 | 1.8 |
| Archaea | Other Archaea | Other Archaea | 7.7 | 7.7 | 7.7 |
| Bacteria | Chloroflexi | Anaerolineaceae | 2.8 | 2.7 | 2.8 |
| Bacteria | Bacteroidota | Bacteroidetes BD2-2 | 1.3 | 1.3 | 1.3 |
| Bacteria | Calditrichota | Calditrichaceae | 0.5 | 0.7 | 0.6 |
| Bacteria | Desulfobacterota | Desulfatiglandaceae | 0.9 | 0.9 | 0.9 |
| Bacteria | Desulfobacterota | Desulfobacteraceae | 1.7 | 2.1 | 1.9 |
| Bacteria | Desulfobacterota | Desulfobulbaceae | 0.6 | 0.8 | 0.7 |
| Bacteria | Desulfobacterota | Desulfocapsaceae | 1.0 | 1.3 | 1.1 |
| Bacteria | Desulfobacterota | Desulfosarcinaceae | 5.2 | 6.5 | 5.8 |
| Bacteria | Bacteroidota | Flavobacteriaceae | 1.9 | 1.7 | 1.8 |
| Bacteria | Proteobacteria | Halieaceae | 0.6 | 0.6 | 0.6 |
| Bacteria | Latescibacterota | Latescibacteraceae | 0.9 | 0.7 | 0.8 |
| Bacteria | Cloacimonadota | MSBL8 | 1.0 | 0.9 | 1.0 |
| Bacteria | Bacteroidota | Marinilabiliaceae | 0.6 | 0.7 | 0.7 |
| Bacteria | Other Bacteria | Other Bacteria | 41.1 | 41.4 | 41.3 |
| Bacteria | Planctomycetota | Pirellulaceae | 1.4 | 1.2 | 1.3 |
| Bacteria | Planctomycetota | SG8-4 | 0.9 | 1.1 | 1.0 |
| Bacteria | Myxococcota | Sandaracinaceae | 1.0 | 1.1 | 1.1 |
| Bacteria | Spirochaetota | Spirochaetaceae | 1.0 | 1.0 | 1.0 |
| Bacteria | Campylobacterota | Sulfurimonadaceae | 0.8 | 0.9 | 0.8 |
| Bacteria | Campylobacterota | Sulfurovaceae | 7.0 | 7.9 | 7.4 |
| Bacteria | Acidobacteriota | Thermoanaerobaculaceae | 1.2 | 0.7 | 0.9 |

**Table S3:** Concentrations and δ^13^C values of fatty acids, alcohols and hydrocarbons from the original cold seep sediment used for incubation. Quantification was based on internal standards of known concentration added prior to extraction. MAGE: monoalkyl glycerol ether. DAGE: dialkyl glycerol ether. OH: hydroxy.

| **Fatty Acids** | | |  | **Alcohols/Hydrocarbons** | | |
| --- | --- | --- | --- | --- | --- | --- |
| **Name** | **µg g_dw_^-1^** | **δ^13^C [‰]** |  | **Name** | **µg g_dw_^-1^** | **δ^13^C [‰]** |
| aiC13:0 | 0.27 | -37 |  | OH-C14:0 | 1.52 | -36 |
| iC13:0 | 0.73 | -40 |  | Crocetane | 1.26 | -100 |
| C13:0 | 0.52 | -31 |  | OH-C16:1ω7c | 2.35 | -78 |
| i14:0 | 1.67 | -32 |  | OH-C16:1ω5c | 2.53 | -77 |
| C14:1ω7 | 0.98 | -28 |  | OH-C16:0 | 1.84 | -57 |
| C14:1ω5 | 0.74 | -36 |  | OH-C18:0 | 0.77 | -29 |
| C14:0 | 9.86 | -38 |  | Phytol | 10.89 | -29 |
| aiC15:0 | 5.29 | -38 |  | PMI | 0.24 | -38 |
| iC15:0 | 7.32 | -38 |  | MAGE C14:0 | 1.24 | -66 |
| C15:0 | 2.56 | -34 |  | OH-C20:0 | 0.29 | -33 |
| iC16:1 | 0.77 | -27 |  | MAGE C15:0 | 0.47 | -24 |
| iC16:0 | 1.5 | -32 |  | MAGE C16:1ω7c | 1.33 | -66 |
| C16:1ω7c | 14.08 | -34 |  | MAGE C16:1ω5c | 1.75 | -63 |
| C16:1ω7t | 3.58 | -40 |  | MAGE C16:0 | 1.46 | -48 |
| C16:1ω5c | 20.71 | -61 |  | OH-C22:0 | 2.45 | -28 |
| C16:0 | 23.4 | -33 |  | Squalene | 0.76 | -36 |
| iC17:1 | 1.00 | -41 |  | MAGE sn2-Phy | 1.23 | -99 |
| 10MeC16:0 | 1.80 | -48 |  | OH-C24:0 | 1.51 | -31 |
| aiC17:0 | 0.71 | -29 |  | OH-C26:0 | 1.14 | -30 |
| iC17:0 | 0.8 | -33 |  | Cholesterol | 5.25 | -26 |
| C17:1ω8c | 0.45 | -30 |  | Sitosterol | 6.97 | -26 |
| C17:1ω6c | 1.59 | -46 |  | Dinosterol | 3.99 | -24 |
| cyC17:0ω5,6 | 1.09 | -59 |  | Tetrahymanol | 0.92 | -29 |
| C17:0 | 0.78 | -33 |  | Diplopterol | 1.38 | -41 |
| C18:1ω9c | 5.77 | -30 |  | DAGE 32:2 | 5.55 | -77 |
| C18:1ω7c | 6.98 | -32 |  | Archaeol | 3.99 | -91 |
| C18:0 | 4.02 | -32 |  | sn2-OH-Archaeol | 7.41 | -101 |
| 2MeC19:0 (IS) | 1.04 | -32 |  |  |  |  |
| C19:0 | 0.58 | -30 |  |  |  |  |
| C20:4 (EPA) | 0.95 | -26 |  |  |  |  |
| C20:5ω3 | 2.02 | -26 |  |  |  |  |
| C20:0 | 0.83 | -31 |  |  |  |  |
| C22:0 | 1.49 | -31 |  |  |  |  |
| C24:0 | 1.79 | -31 |  |  |  |  |
| C26:0 | 0.97 | -32 |  |  |  |  |
| C28:0 | 0.52 | -34 |  |  |  |  |

**Table S4:** Shift of δ^13^C values (Δδ^13^C, ‰) of lipid biomarkers characteristic for the AOM performing microorganisms in all treatments with the different ^13^C-labeled and non-labeled carbon sources. FA: fatty acid. MAGE: monoalkyl glycerol ether. DAGE: dialkyl glycerol ether. OH: Hydroxy.

|  |  | **Δδ13C [‰]** | | | | | | | | | | | |
| --- | --- | --- | --- | --- | --- | --- | --- | --- | --- | --- | --- | --- | --- |
|  |  | **Treatment** | **DIC w/ ^13^CH_4_** | | | **DI^13^C w/ CH_4_** | | | **DI^13^C w/o CH_4_** | | | **DIC w/ ^13^CH_4_** | **DI^13^C w/ CH_4_** |
|  |  | **Time [days]** | **10** | **20** | **30** | **10** | **20** | **30** | **10** | **20** | **30** | **Control (30)** | **Control (30)** |
| **Archaeal lipids** |  | **Crocetane** | 6.3 | 17.8 | 36.9 | 9.6 | 65.6 | 126 | 0.3 | 5.3 | 1.6 | 0.3 | 0.3 |
|  |  | **sn2 phy MAGE** | 6.9 | 16.1 | 26.1 | 7.9 | 45.4 | 78.4 | 4.6 | 10.4 | 10.2 | 0.8 | 0.8 |
|  |  | **Archaeol** | 8.7 | 19.1 | 27.7 | 10.5 | 39.1 | 71 | 1.6 | 5.3 | 7.4 | 0.7 | 0.2 |
|  |  | **sn2 OH Archaeol** | 2.2 | 10.6 | 13.6 | 3.4 | 12.8 | 31.8 | 0.9 | 2.7 | 8.9 | 0.9 | 0.9 |
| **Bacterial lipids** | **Fatty acids (FA)** | **C14:0** | 2.3 | 17.2 | 30.1 | 34 | 133 | 172 | 2.8 | 6 | 7.3 | 0.4 | 0.4 |
|  |  | **iC15:0** | 0.8 | 0.8 | 1.3 | 8.3 | 18.1 | 28.8 | 7.8 | 9.2 | 14.1 | 0.3 | 0.8 |
|  |  | **aiC15:0** | 0.8 | 3.1 | 5 | 12.5 | 26.6 | 37.8 | 8.8 | 15.8 | 21.3 | 0.7 | 0.3 |
|  |  | **C16:1ω7c** | 1.1 | 20.1 | 30 | 39.1 | 131 | 197 | 7.8 | 11.2 | 11.3 | 1.6 | 0.7 |
|  |  | **C16:1ω5c** | 6 | 35.6 | 55.6 | 72.8 | 268 | 393 | 11.1 | 13.5 | 13.5 | 0.3 | 0.7 |
|  |  | **C16:0** | 1.2 | 12 | 19.7 | 30.3 | 95.7 | 129 | 3.5 | 5.9 | 7 | 1.2 | 0.7 |
|  |  | **cyC17:0ω5,6** | 5.7 | 22.2 | 32.8 | 31.2 | 173 | 258 | 5.7 | 5.2 | 6.6 | 0.5 | 0.5 |
|  | **Alcohols (OH)** | **C18:1ω7c** | 0 | 0.5 | 3.1 | 10.9 | 21.8 | 37.9 | 8.1 | 9.9 | 10.4 | 0 | 0 |
|  |  | **C16:1ω7c** | 1.7 | 5.6 | 7.9 | 1 | 24.9 | 36 | 0.4 | 0.6 | 0.7 | 2.3 | 1.5 |
|  |  | **C16:1ω5c** | 0 | 6.6 | 11.6 | 8 | 26.7 | 45 | 0.2 | 1.8 | 1.9 | 1.8 | 2 |
|  |  | **C16:0** | 1.1 | 1.4 | 3 | 0.8 | 5.4 | 8.2 | 0.5 | 0.3 | 2.1 | 2 | 2.3 |
|  | **MAGEs** | **C16:1ω7c** | 6.1 | 15.1 | 24.2 | 14.3 | 110 | 190 | 1.3 | 1.2 | 2.5 | 2.6 | 0.6 |
|  |  | **C16:1ω5** | 4.6 | 4.9 | 7.5 | 8.7 | 38.2 | 62 | 2.5 | 3.3 | 3 | 1.8 | 2.1 |
|  |  | **C16:0** | 8.2 | 11.2 | 14.3 | 15.4 | 35.1 | 59 | 2.6 | 4.2 | 4.4 | 2.8 | 2.8 |
|  | **DAGE** | **C32:2** | 3.6 | 9 | 13.1 | 1.5 | 26 | 60.4 | 1.6 | 4.8 | 3.3 | 2.5 | 0.2 |

**Table S5:** Carbon assimilation rates in µg C g_dw_^-1^ y^-1^ of characteristic lipid biomarkers calculated with DIC as ^13^C source for all treatments. FA: fatty acid. MAGE: monoalkyl glycerol ether. DAGE: dialkyl glycerol ether. OH: Hydroxy.

|  |  | **Inorganic carbon ^13^C carbon assimilation [µg C g_dw_^-1^ y^-1^]** | | | | | | | | | | | |
| --- | --- | --- | --- | --- | --- | --- | --- | --- | --- | --- | --- | --- | --- |
|  |  | **Treatment** | **DIC w/ ^13^CH_4_** | | | **DI^13^C w/ CH_4_** | | | **DI^13^C w/o CH_4_** | | | **DIC w/ ^13^CH_4_** | **DI^13^C w/ CH_4_** |
|  |  | **Type** | **Living sample** | | | | | | | | | **Killed control** | |
|  |  | **Time [days]** | **10** | **20** | **30** | **10** | **20** | **30** | **10** | **20** | **30** | **30** | **30** |
| **Archaeal lipids** |  | **Crocetane** | 0.224 | 0.241 | 0.313 | 0.066 | 0.230 | 0.299 | 0.002 | 0.016 | 0.004 | 0.003 | 0.001 |
|  |  | **sn2 phy MAGE** | 0.214 | 0.189 | 0.191 | 0.047 | 0.137 | 0.160 | 0.026 | 0.028 | 0.020 | 0.008 | 0.002 |
|  |  | **Archaeol** | 0.917 | 0.765 | 0.691 | 0.213 | 0.404 | 0.494 | 0.031 | 0.049 | 0.051 | 0.026 | 0.002 |
|  |  | **sn2 OH Archaeol** | 0.425 | 0.766 | 0.616 | 0.124 | 0.239 | 0.401 | 0.032 | 0.044 | 0.110 | 0.063 | 0.011 |
| **Bacterial lipids** | **Fatty acids (FA)** | **C14:0** | 0.605 | 1.71 | 1.86 | 1.70 | 3.41 | 2.97 | 0.132 | 0.138 | 0.123 | 0.036 | 0.008 |
|  |  | **iC15:0** | 0.103 | 0.042 | 0.039 | 0.209 | 0.233 | 0.250 | 0.187 | 0.106 | 0.119 | 0.015 | 0.007 |
|  |  | **aiC15:0** | 0.142 | 0.216 | 0.215 | 0.436 | 0.473 | 0.454 | 0.290 | 0.251 | 0.250 | 0.045 | 0.004 |
|  |  | **C16:1ω7c** | 0.392 | 2.72 | 2.52 | 2.67 | 4.56 | 4.62 | 0.505 | 0.348 | 0.259 | 0.203 | 0.017 |
|  |  | **C16:1ω5c** | 3.127 | 7.08 | 6.88 | 7.30 | 13.8 | 13.6 | 1.06 | 0.618 | 0.456 | 0.060 | 0.025 |
|  |  | **C16:0** | 0.725 | 2.85 | 2.92 | 3.63 | 5.87 | 5.316 | 0.400 | 0.321 | 0.281 | 0.263 | 0.028 |
|  |  | **cyC17:0ω5,6** | 0.159 | 0.235 | 0.217 | 0.167 | 0.473 | 0.476 | 0.029 | 0.013 | 0.012 | 0.005 | 0.001 |
|  | **Alcohols (OH)** | **C18:1ω7c** | 0.000 | 0.032 | 0.130 | 0.373 | 0.381 | 0.447 | 0.262 | 0.155 | 0.120 | 0.000 | 0.000 |
|  |  | **C16:1ω7c** | 0.104 | 0.133 | 0.117 | 0.012 | 0.153 | 0.149 | 0.004 | 0.003 | 0.003 | 0.053 | 0.006 |
|  |  | **C16:1ω5c** | 0.003 | 0.169 | 0.185 | 0.104 | 0.177 | 0.201 | 0.002 | 0.011 | 0.008 | 0.044 | 0.009 |
|  |  | **C16:0** | 0.055 | 0.026 | 0.034 | 0.008 | 0.026 | 0.026 | 0.005 | 0.001 | 0.007 | 0.035 | 0.007 |
|  | **MAGEs** | **C16:1ω7c** | 0.200 | 0.187 | 0.186 | 0.089 | 0.350 | 0.409 | 0.008 | 0.003 | 0.005 | 0.030 | 0.001 |
|  |  | **C16:1ω5** | 0.198 | 0.079 | 0.076 | 0.071 | 0.161 | 0.176 | 0.019 | 0.012 | 0.008 | 0.028 | 0.006 |
|  |  | **C16:0** | 0.290 | 0.149 | 0.119 | 0.104 | 0.121 | 0.137 | 0.017 | 0.013 | 0.010 | 0.036 | 0.006 |
|  | **DAGE** | **C32:2** | 0.539 | 0.509 | 0.463 | 0.042 | 0.380 | 0.595 | 0.044 | 0.063 | 0.032 | 0.137 | 0.002 |

**Table S6:** Carbon assimilation rates in µg C g_dw_^-1^ y^-1^ of lipid biomarkers characteristic for AOM communities calculated with CH_4_ as ^13^C source. Values for bacterial lipids are hypothetical as SRB are autotrophs. FA: fatty acid. MAGE: monoalkyl glycerol ether. DAGE: dialkyl glycerol ether. OH: Hydroxy.

|  |  | **Methane carbon ^13^C carbon assimilation [µg C g_dw_^-1^ y^-1^]** | | | | |
| --- | --- | --- | --- | --- | --- | --- |
|  |  | **Treatment** | **DIC w/ ^13^CH_4_** | | | **DIC w/ ^13^CH_4_** |
|  |  |  | **Living sample** | | | **Killed control** |
|  |  | **Time [days]** | **10** | **20** | **30** | **30** |
| **Archaeal lipids** |  | **Crocetane** | 0.026 | 0.025 | 0.044 | 0.026 |
|  |  | **MAGE sn2 phy** | 0.025 | 0.020 | 0.027 | 0.025 |
|  |  | **Archaeol** | 0.107 | 0.081 | 0.098 | 0.107 |
|  |  | **sn2 OH Archaeol** | 0.050 | 0.081 | 0.087 | 0.050 |
| **Bacterial lipids** | **Fatty acids (FA)** | **C14:0** | 0.012 | 0.004 | 0.006 | 0.012 |
|  |  | **iC15:0** | 0.071 | 0.180 | 0.263 | 0.071 |
|  |  | **aiC15:0** | 0.017 | 0.023 | 0.031 | 0.017 |
|  |  | **C16:1ω7c** | 0.023 | 0.008 | 0.011 | 0.023 |
|  |  | **C16:1ω5c** | 0.367 | 0.745 | 0.977 | 0.367 |
|  |  | **C16:0** | 0.085 | 0.299 | 0.414 | 0.085 |
|  |  | **cyC17:0ω5,6** | 0.019 | 0.025 | 0.031 | 0.019 |
|  | **Alcohols (OH)** | **C18:1ω7c** | 0.000 | 0.003 | 0.018 | 0.000 |
|  |  | **C16:1ω7c** | 0.012 | 0.014 | 0.017 | 0.012 |
|  |  | **C16:1ω5c** | 0.000 | 0.018 | 0.026 | 0.000 |
|  |  | **C16:0** | 0.006 | 0.003 | 0.005 | 0.006 |
|  | **MAGEs** | **C16:1ω7c** | 0.023 | 0.020 | 0.026 | 0.023 |
|  |  | **C16:1ω5** | 0.046 | 0.286 | 0.358 | 0.046 |
|  |  | **C16:0** | 0.034 | 0.016 | 0.017 | 0.034 |
|  | **DAGE** | **C32:2** | 0.063 | 0.054 | 0.066 | 0.063 |

**Table S7:** Calculated lipid turnover times in years of characteristic lipid biomarkers calculated with DIC as ^13^C source for all treatments. FA: fatty acid. MAGE: monoalkyl glycerol ether. DAGE: dialkyl glycerol ether. OH: Hydroxy.

|  |  | **Lipid turnover time [years] – inorganic carbon source** | | | | | | | | | |
| --- | --- | --- | --- | --- | --- | --- | --- | --- | --- | --- | --- |
|  |  | **Treatment** | **DIC w/ ^13^CH_4_** | | | **DI^13^C w/ CH_4_** | | | **DI^13^C w/o CH_4_** | | |
|  |  | **Time [days]** | **10** | **20** | **30** | **10** | **20** | **30** | **10** | **20** | **30** |
| **Archaeal lipids** |  | **Crocetane** | 4.8 | 4.5 | 3.5 | 16.5 | 4.7 | 3.6 | 665 | 66.1 | 297 |
|  |  | **MAGE sn2 phy** | 4.4 | 4.9 | 4.9 | 20.0 | 6.8 | 5.9 | 36.1 | 33.4 | 45.9 |
|  |  | **Archaeol** | 3.5 | 4.2 | 4.6 | 15.1 | 7.9 | 6.5 | 103 | 65.5 | 63.1 |
|  |  | **sn2 OH Archaeol** | 13.6 | 7.6 | 9.4 | 46.7 | 24.3 | 14.5 | 184 | 131 | 52.5 |
| **Bacterial lipids** | **Fatty acids (FA)** | **C14:0** | 13.1 | 4.6 | 4.3 | 4.6 | 2.3 | 2.7 | 59.8 | 57.4 | 64.5 |
|  |  | **iC15:0** | 38.7 | 94.7 | 103 | 19.1 | 17.1 | 16.0 | 21.3 | 37.6 | 33.4 |
|  |  | **aiC15:0** | 38.7 | 25.6 | 25.7 | 12.7 | 11.7 | 12.1 | 19.0 | 22.0 | 22.1 |
|  |  | **C16:1ω7c** | 27.5 | 4.0 | 4.3 | 4.0 | 2.4 | 2.3 | 21.4 | 31.0 | 41.7 |
|  |  | **C16:1ω5c** | 5.1 | 2.2 | 2.3 | 2.2 | 1.2 | 1.2 | 14.9 | 25.7 | 34.8 |
|  |  | **C16:0** | 26.2 | 6.7 | 6.5 | 5.2 | 3.2 | 3.6 | 47.4 | 59.1 | 67.5 |
|  |  | **cyC17:0ω5,6** | 5.3 | 3.6 | 3.9 | 5.1 | 1.8 | 1.8 | 29.3 | 66.8 | 71.1 |
|  | **Alcohols (OH)** | **C18:1ω7c** | NA | 169 | 41.7 | 14.5 | 14.2 | 12.1 | 20.6 | 34.9 | 45.1 |
|  |  | **C16:1ω7c** | 18.2 | 14.3 | 16.3 | 155 | 12.4 | 12.8 | 475 | 598 | 691 |
|  |  | **C16:1ω5c** | 756 | 12.1 | 11.1 | 19.7 | 11.6 | 10.2 | 875 | 189 | 242 |
|  |  | **C16:0** | 26.8 | 57.0 | 43.3 | 188 | 56.9 | 56.2 | 308 | 1387 | 226 |
|  | **MAGEs** | **C16:1ω7c** | 5.0 | 5.3 | 5.3 | 11.1 | 2.8 | 2.4 | 131 | 287 | 185 |
|  |  | **C16:1ω5** | 6.6 | 16.4 | 17.2 | 18.2 | 8.1 | 7.4 | 67.8 | 105 | 158 |
|  |  | **C16:0** | 3.7 | 7.1 | 9.0 | 10.3 | 8.8 | 7.8 | 63.7 | 82.4 | 107 |
|  | **DAGE** | **C32:2** | 8.4 | 8.9 | 9.8 | 109 | 11.9 | 7.6 | 103 | 71.7 | 142 |

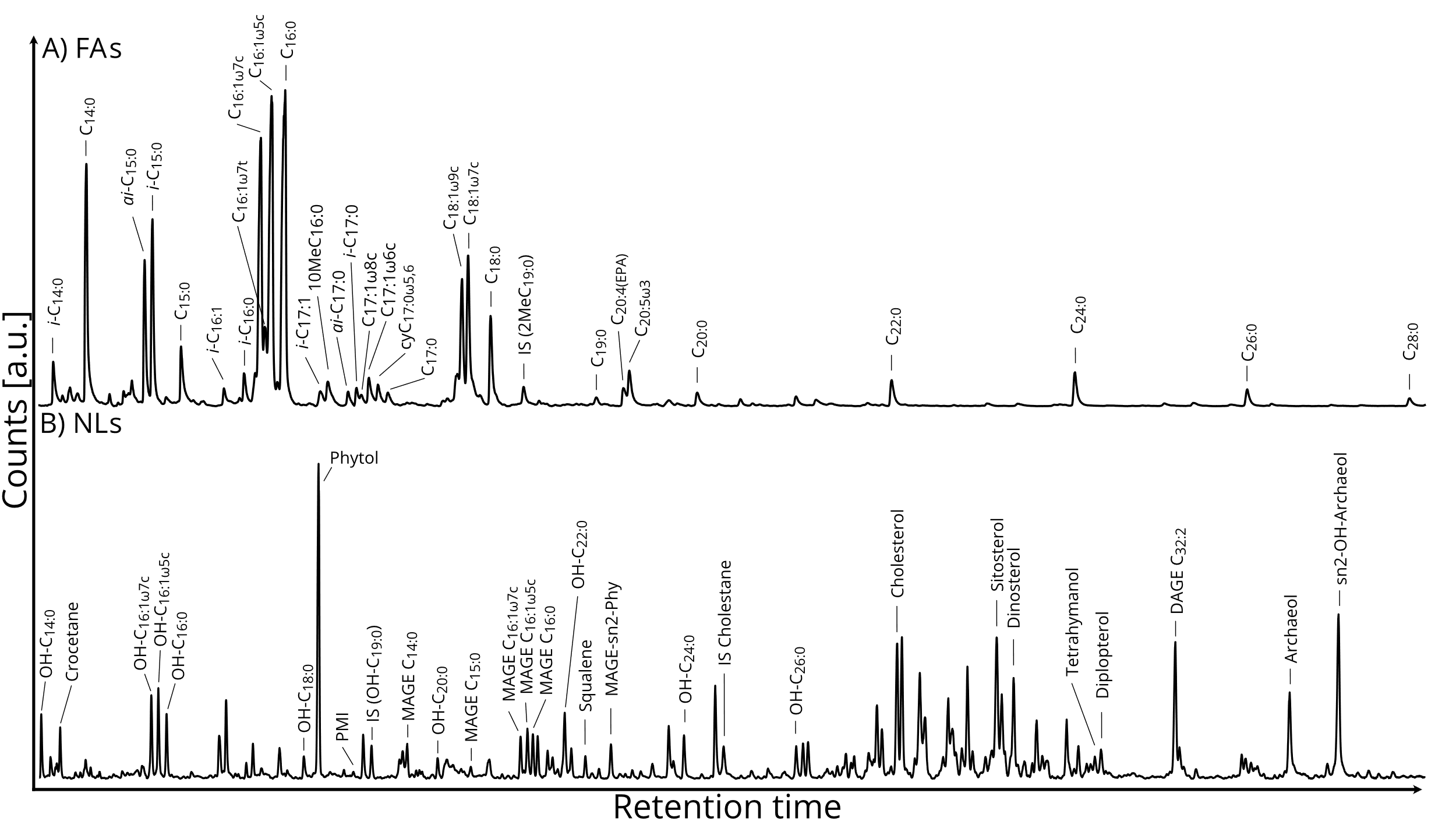

**Figure S1:** Partial gas chromatograms of **(A)** the fatty acid (FA) fraction (analyzed as fatty acid methyl esters) and **(B)** the neutral lipid (NL) fraction, containing alcohols as trimethylsilyl derivatives and hydrocarbons, obtained from the original sediment material used for the SIP experiments.

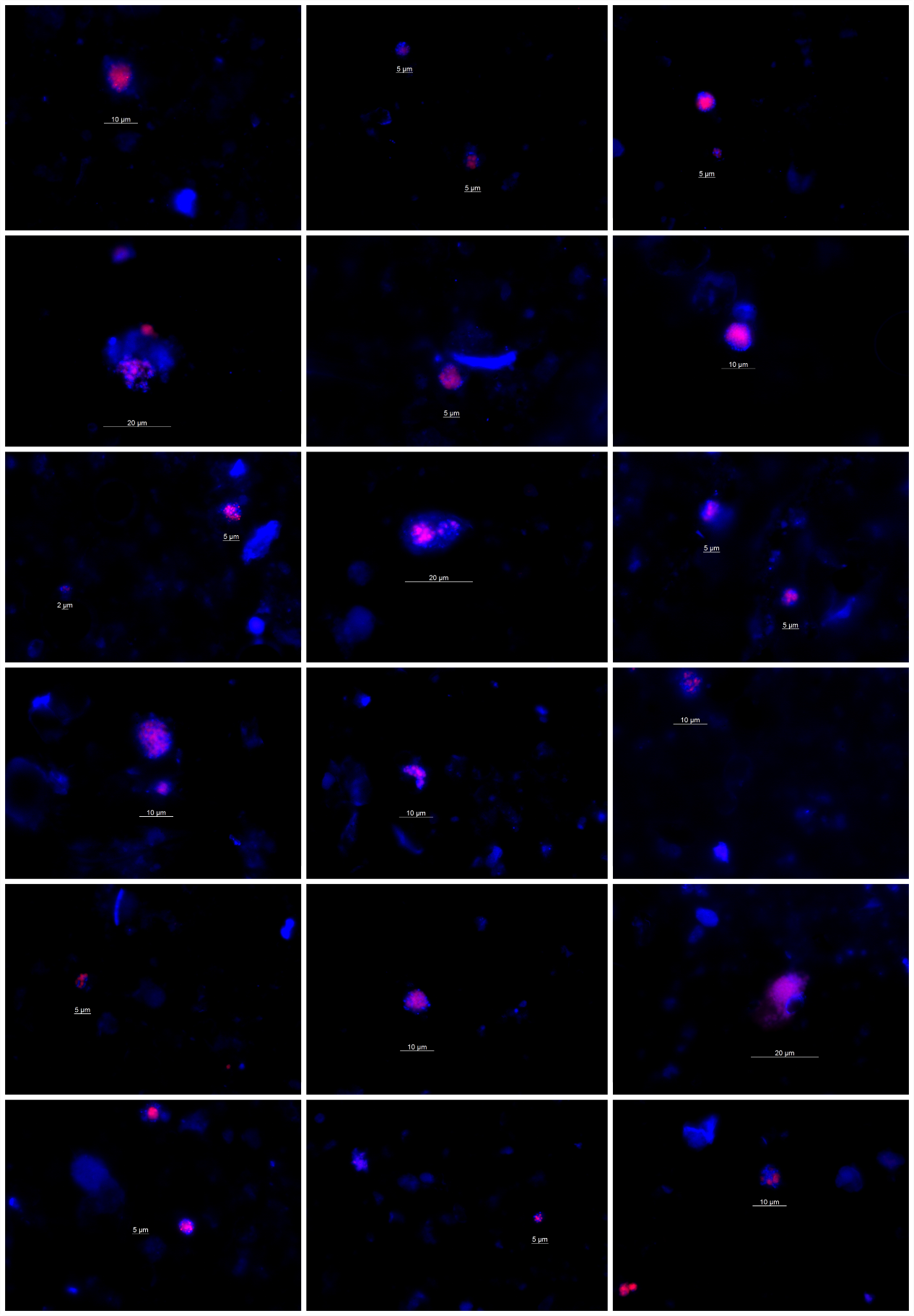

**Figure S2:** Selection of fluorescence micrographs of the natural AOM enrichment at the start (0 days) and end (30 days) of the incubations. Each image depicts DAPI stained DNA (all cells, blue) and ANME-2 specific probe-stained archaeal cells (red).
